# A practical guide to time-resolved fluorescence microscopy and spectroscopy

**DOI:** 10.1101/2024.01.25.577300

**Authors:** Benjamin S. Clark, Irene Silvernail, Kenya Gordon, Jose F. Castaneda, Andi N. Morgan, Lewis A. Rolband, Sharonda J. LeBlanc

## Abstract

Time-correlated single photon counting (TCSPC) coupled with confocal microscopy is a versatile biophysical tool that enables real-time monitoring of biomolecular dynamics across many timescales. With TCSPC, Fluorescence correlation spectroscopy (FCS) and pulsed interleaved excitation-Förster resonance energy transfer (PIE-FRET) are collected simultaneously on diffusing molecules to extract diffusion characteristics and proximity information. This article is a guide to calibrating FCS and PIE-FRET measurements with several biological samples including liposomes, streptavidin-coated quantum dots, proteins, and nucleic acids for reliable determination of diffusion coefficients and FRET efficiency. The FRET efficiency results are also compared to surface-attached single molecules using fluorescence lifetime imaging microscopy (FLIM-FRET). Combining the methods is a powerful approach to revealing mechanistic details of biological processes and pathways.

## INTRODUCTION

Fluorescence microscopy is an indispensable tool for biological and materials research. Confocal fluorescence microscopes that produce diffraction-limited images with single-molecule resolution are commonly available in laboratories and microscopy cores worldwide. These advances have had an immeasurable impact on many fields, from cell biology to materials science. Super-resolution microscopy, which earned the 2014 Nobel Prize in Chemistry, enabled sub-diffraction limited optical imaging and spatial resolution down to a few nanometers (1). Such length scales are highly relevant for molecular and cellular biology, and one can resolve sub-cellular structures with exquisite detail.

While high spatial resolution optical images yield the localization and dynamics of biomolecules in living cells and organisms, time-resolved fluorescence methods add several experimental dimensions. Traditional confocal microscopy solely utilizes the intensity of fluorescent molecules to report location. In addition to intensity, fluorescent molecules possess another intrinsic photophysical property called the excited state lifetime or ‘fluorescence lifetime’. An excited state is created in a molecule upon activation by a suitable laser pulse that promotes ground state electrons to a higher energy level. The average time between this excitation and subsequent relaxation by photon emission is the fluorescence lifetime. Fluorophores such as organic dyes, quantum dots, and fluorescent proteins have distinct fluorescence lifetimes, which may be distinguished with time-resolved measurements. Fluorescence lifetime imaging microscopy (FLIM) (2,3) also reports on the local environment in live cells or of surface-attached molecules. These quantitative optical images combine the high-resolution fluorescence intensity and lifetime measurements into a single image. In addition, a droplet containing freely diffusing molecules can also be probed, and one can produce time-resolved spectroscopic data by analyzing photon bursts as fluorophores traverse the confocal volume.

Obtaining time-resolved fluorescence data with high resolution requires specialized photon-counting equipment in addition to the standard confocal microscope components. Time-correlated single photon counting (TCSPC) (4,5) collects photon data that enables simultaneous fluorescence intensity, lifetime, and correlation measurements. Spatially and spectrally resolved TCSPC captures dynamic processes across time scales from sub-nanoseconds to seconds. These tools are applied to directly investigate excited state dynamics in nanoscale optical materials (6,7), or in biomolecules to observe the transient interactions and conformational dynamics that drive biology (8-10). The time-tagged data is spatially mapped to rapidly produce FLIM images for samples that adhere to a surface. TCSPC measurements for freely diffusing molecules in a solution require focusing a laser beam several micrometers above a microscope coverslip into the droplet solution (Figure 1A). Photons are detected from a fixed location within the droplet (as opposed to scanning the coverslip surface to acquire an image). The fluorescence intensity traces collected from single molecules diffusing through the femtoliter confocal volume are analyzed by fluorescence correlation spectroscopy (FCS) (11-14) or fluorescence cross correlation spectroscopy (FCCS) (15,16) to quantitate diffusion characteristics, concentration, and stoichiometry of the diffusing species. More advanced correlation analysis of the time-tagged photon data is also possible, such as fluorescence blinking (7) and photon antibunching (6).

**Figure 1:**
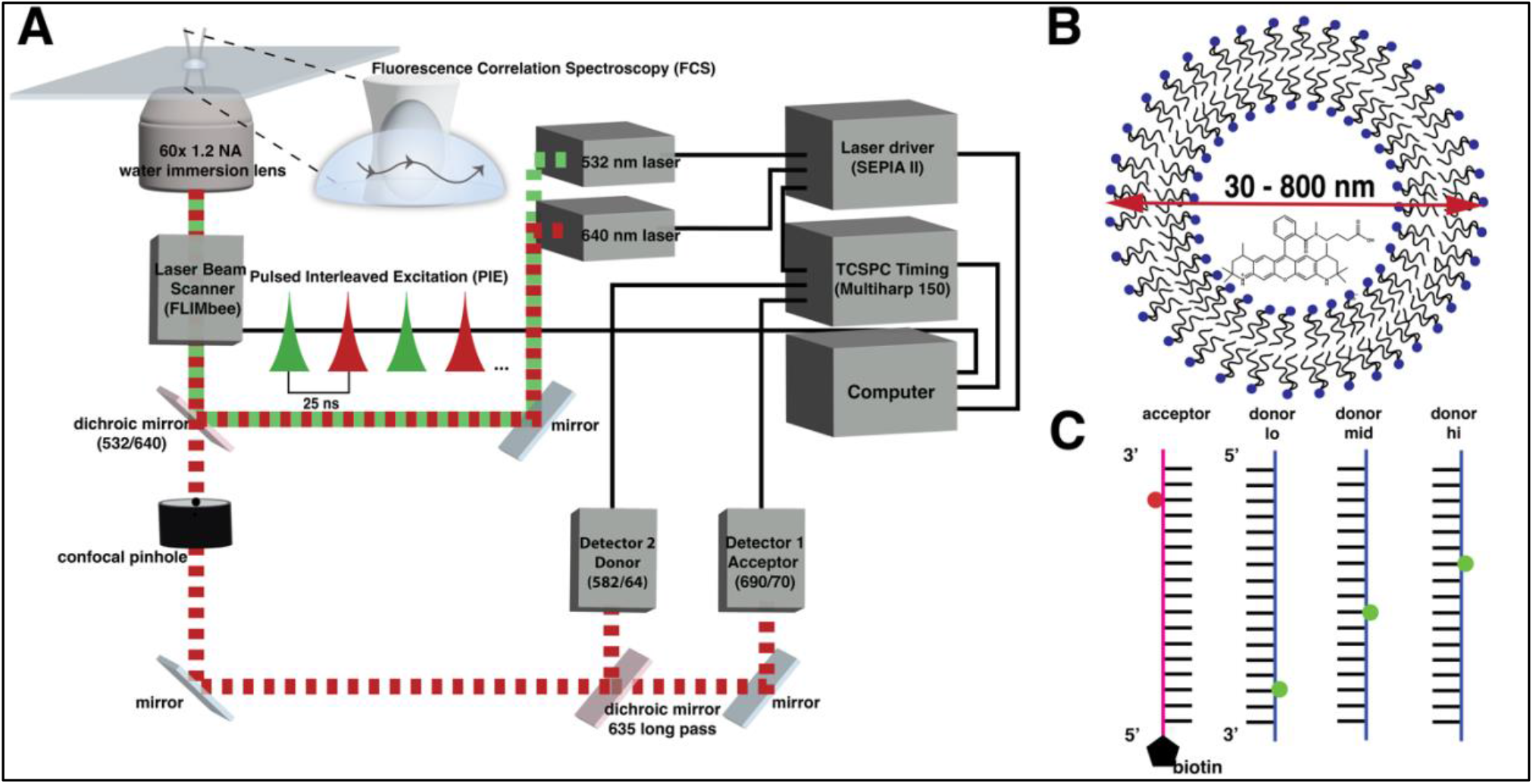
**A)** Time-resolved confocal fluorescence microscope schematic. A droplet sample is placed on a thin coverslip, and the is laser focused several micrometers into the droplet for diffusing measurements such as FCS and PIE-FRET. Two diode lasers are driven in pulsed interleaved excitation mode and excite the sample through a high numerical aperture objective lens. Fast laser beam scanning is used to image surface-attached molecules. Donor and acceptor molecules are excited sequentially on the nanosecond timescale, the fluorescence is passed through a dichroic mirror and confocal pinhole and detected by single photon counting modules. Each photon is time-tagged by the TCSPC event timer. **B)** Liposome encapsulation strategy. The fluorescent dye Atto550 was encapsulated in liposomes of varying sizes (30 nm – 800 nm) and the fluorescent spherical particles were used to calibrate FCS measurements. **C)** Schematic of benchmark DNA oligomers for PIE-FRET calibration. An acceptor DNA strand containing a 5’ biotin was fluorescently labeled near the 3’ end. The complementary DNA strands were labeled at sites 11, 15, and 23 bases away from the acceptor label. When annealed, lo, mid and hi FRET benchmark samples are produced. The sequences were outlined in an smFRET benchmark study (51) and are listed in Supplementary Table S1.

The applications of time-resolved fluorescence measurements for biophysical studies of both surface-attached and freely diffusing biomolecules are numerous. FCS and its variations (15,17-19) have been utilized to measure concentrations and diffusion coefficients of biomolecules *in vitro* and in living cells (19-21), but the measurement is limited to diffusing species. A complimentary technique (22) is Förster/fluorescence resonance energy transfer (FRET) or single-molecule FRET (smFRET), which can be observed on surface-attached or diffusing molecules (23). Attaching donor and acceptor reporters to biomolecules is a robust tool to study individual protein and nucleic acid conformational dynamics (9,24-37). Adding to its versatility, FRET is detected by measuring changes in intensity or changes in the fluorescence lifetime of the donor molecule. The phenomenon may be combined with other measurements such as FCS (22,29,38), FLIM (3,39,40), fluorescence lifetime correlation spectroscopy (FLCS) (41,42), and super-resolution microscopy (43,44). Measuring FRET between single molecules (smFRET) was advanced by rapidly alternating two laser pulses on the micro/millisecond timescale (alternating laser excitation - ALEX) (45,46) or nanosecond timescale (pulsed interleaved excitation - PIE) (47-50) to excite the donor and acceptor molecules sequentially and detect the fluorescence signals quasi-simultaneously. The need to validate results obtained from smFRET measurements on different types of microscopes in labs worldwide has led to several important benchmark studies (10,51,52) that formulate a set of community standards. Calculating relative changes in apparent FRET efficiency can be sufficient to investigate transient biomolecular interactions or conformational dynamics. In contrast, for smFRET-aided structural modeling (53-56), one needs to recover the true FRET efficiency and convert it to a physical distance, which must be carefully approached.

Here, we compare methods to calibrate the diffusion coefficient (using FCS), FRET efficiency, and fluorescence lifetime for several biomolecular samples using our commercially available PicoQuant MicroTime200 microscope. The collected photon data spans from sub-nanosecond to seconds. Each type of data requires specialized analysis that if properly validated can simultaneously reveal conformational dynamics, interactions, and oligomeric states of a diffusing biomolecular system. We fluorescently labeled a set of standard DNA samples purchased with sequences from a previous smFRET benchmark study (51) and simultaneously measured FCS, PIE-FRET, and fluorescence lifetimes on the diffusing molecules. We also measured a hybrid RNA/DNA substrate. FCS diffusion coefficients were validated by encapsulating a fluorescent dye in liposomes of different sizes (Figure 1B), measuring the diffusion coefficients of the liposomes and several approximately spherical particles, and fitting the results to the theoretical Stokes-Einstein equation (57).

To calibrate our diffusing PIE-FRET measurements, the average FRET efficiency for annealed DNA samples (Figure 1C) was calculated using an intensity and lifetime-based approach, and the results were compared to the benchmark study (51). The intensity-based FRET calculation requires determining three correction factors (*α, δ*, and *γ*), for which we have presented two methods. We also compared the calculated FRET efficiency of the same benchmark DNA samples when surface-attached to a microscope coverslip by obtaining FLIM-FRET images. The FRET efficiency for individual surface-attached DNA duplexes was calculated using the intensity and lifetime data. Our side-by-side comparisons of several methods and analysis approaches demonstrate good agreement, which highlights the robustness of single-molecule fluorescence approaches when community standards are followed.

## MATERIALS AND METHODS

### Time-resolved fluorescence microscopy and spectroscopy

Time-resolved fluorescence measurements were collected using a custom MicroTime 200 (PicoQuant - Berlin, Germany) with SymPhoTime64 software for data acquisition and analysis (Figure 1A). The modular time-resolved confocal microscope system is built around an inverted Olympus IX-83 microscope body with a side port for optics and a piezoelectric z stage. It is equipped with three picosecond pulsed diode lasers (485, 531, and 636 nm) and a laser driver module (SEPIA II) capable of operating in pulsed (up to 80 MHz) and continuous wave (cw) modes. The lasers are cleaned up with narrowband filters, fiber-coupled into the main optical unit (MOU), directed to a main quad-band dichroic mirror, and focused onto the sample with a 60x 1.2 numerical aperture (NA) water immersion lens (Olympus UPlanSApo, Superachromat). A fast galvo beam scanning module (FLIMbee) with a 0.5 μs dwell time (lower limit) is used for laser beam scanning.

Fluorescence from the sample is collected with the same objective, and spatially filtered through an exchangeable circular confocal pinhole (100 μm used for these experiments) and directed to one or two single photon avalanche diodes (SPADs – Excelitas). For Förster/fluorescence resonance energy transfer (FRET) experiments with 531 and 636 nm laser excitation, the fluorescence is spectrally filtered using a dichroic mirror (635 long pass) placed between the two detectors and bandpass filters in front of each detector to separate donor (582/64, Channel 2) and acceptor (690/70, Channel 1) emission. For TCSPC, a multichannel event timer with 10 ps resolution and 650 ps deadtime (MultiHarp 150) in time-tagged time-resolved (TTTR) measurement mode applies time tags to individual photons detected at each SPAD, both relative to the laser pulse (nanotime) and relative to the start of the experiment (macrotime). Generating real-time histograms of the different time tags enables simultaneous fluorescence intensity, lifetime, and correlation data collection. Detected photons are also marked with a location in the beam scan of the sample to enable spatial mapping for FLIM. All fluorescence experiments were performed at room temperature.

Operating the lasers in pulsed interleaved excitation (PIE) mode enables selective excitation and detection of separate donor, acceptor, and FRET signals on the nanosecond (ns) time scale (47). The laser excitation is synchronized in the SymPhoTime software such that the 531 and 636 nm lasers pulse in an alternating manner, successively exciting acceptor and donor molecules directly. Detected single photons are associated with the exciting laser pulse and the arrival detector to separate donor, acceptor, and FRET signals.

### Fluorescent Dye Encapsulation

Fluorescent dyes were encapsulated in small/large unilamellar lipid vesicles (or liposomes) (58-60) to calibrate the FCS measurements. A solution of 1% biotinyl cap-phosphoethanolamine (Avanti Polar Lipids, 870277P) in Egg PC (Avanti Polar Lipids, 840051C) was prepared in chloroform. The lipids were dried in a glass culture tube with an argon gas stream to evaporate the chloroform and then placed in a vacuum chamber until completely dry for several hours to overnight. Dried lipids were rehydrated by gentle vortex in a buffer containing 20 mM Tris HCl, 5 mM MgCl_2_, 100 mM sodium acetate, 2% glucose, and 0.02% cyclooctatetraene for imaging. The addition of glucose is for a glucose oxidase/catalase oxygen scavenging system, which was not used in this study. Atto550 dye (Sigma-Aldrich, 42424) was added to the lipid suspension to a final concentration of 5 nM. The lipid-dye suspension was passed 21 times through an Avanti Polar Lipids mini extruder assembled with various Whatman Nucleopore Track-Etch Membranes: 30, 50, 100, 200, 400, and 800 nanometer (nm) diameter pores. A gravity Sepharose CL-4B (Sigma-Aldrich, 61970-08-9) column was used to separate free dye from the liposomes. The resulting liposomes were expected to be spherical with a diameter close to the membrane pore size (61) with encapsulated Atto550 dye (Figure 1B).

### Protein expression and purification

#### SARS-CoV-2 Nonstructural protein-15 (Nsp15)

SARS-CoV-2 Nsp15 WT and mutant plasmids were created as described in prior work by our collaborators (62). A 20 mL starter culture of Terrific Broth (TB) containing 1% glucose and 100 μg/mL ampicillin was inoculated with transformed *Escherichia coli (E*.*coli)* BL21-Gold(DE3) competent cells and grown overnight at 37°C and 210 rpm. The overnight growth was transferred to an 800 mL culture at 37°C and 210 rpm and grown to an optical density (OD) of 0.9. The temperature was reduced to 16°C, the cells were induced with 0.5 mM isopropyl β-D-1-thiogalactopyranoside (IPTG) and allowed to overexpress Nsp15 overnight. Cells were harvested after a ∼ 16-hour expression and stored at -80°C until purification. To purify Nsp15, cells were resuspended in lysis buffer (50 mM Tris pH 8.0, 500 mM NaCl, 5% glycerol, 5 mM βME, 5 mM imidazole) and Roche cOmplete, EDTA-free protease inhibitor tablets (Sigma-Aldrich, 11873580001) and sonicated on ice for a total time of 6.5 minutes. The cleared lysate was recovered via centrifugation at 11000 rpm at 4°C. 1 mL per milligram protein of cobalt-charged beads (TALON) for affinity chromatography was equilibrated and incubated with the cleared lysate for 45 min at 4°C. The solution was poured into a gravity column, and the beads were washed with wash buffer (50 mM Tris pH 8.0, 500 mM NaCl, 5% glycerol, 50 mM imidazole). Elution buffer (50 mM Tris pH 8.0, 500 mM NaCl, 5% glycerol, 5 mM βME, 250 mM imidazole) was used to elute his-tagged Nsp15 from the TALON resin. Nsp15 was then buffer exchanged with a PD-10 gravity column (Cytiva 17-0851-01) into a low salt buffer (50 mM Tris pH 7.2, 150 mM NaCl, 5% glycerol) before thrombin cleavage. Nsp15 in low salt buffer was supplemented with 2mM βME, 2mM CaCl_2_, and 25 units of thrombin per 1 L of TB used in culture. The cleavage reaction was incubated for 4 hours and then passed through TALON resin to collect his-tag free Nsp15. A Superdex-200 column equilibrated in size exclusion chromatography (SEC) buffer (20 mM HEPES pH 7.5, 150 mM NaCl) was used to isolate active Nsp15 hexamers. SDS-PAGE gels confirmed the presence of Nsp15 (MW_monomer_ ∼ 42 kDa). Purified protein was fluorescently labeled by incubating with a 10x molar excess of maleimide dye for 4 hours to overnight, followed by gravity size exclusion chromatography to separate free dye from labeled protein.

#### Thermus aquaticus MutL

Thermus aquaticus (*Taq*) MutL proteins (with a 6x histidine tag and a single cysteine variation for fluorescent labeling) were expressed in *E*.*coli* BL21(DE3) cells with ampicillin selection and 1 mM IPTG induction at OD 0.7. Cell pellets were disrupted by tip ultrasonication, followed by cell lysate purification with cobalt-charged (TALON) affinity chromatography. Purified *Taq* MutL proteins were labeled with cysteine-maleimide attachment chemistry. Approximately one nanomole of purified protein (250 μL) was pipetted into a sterile 1.5 mL tube and 1 μL of 0.05 mM TCEP was added to reduce disulfide bonds for about 15 minutes. A dried maleimide dye aliquot (20 nanomoles) was dissolved in 1 μL of DMSO. The protein suspension was added to the dissolved dye, and the mixture reacted for 30 minutes at room temperature, followed by 2 hours at cold room conditions. Free dye was removed by purification with a P6 (Bio-Rad) gravity column. SDS-PAGE gels confirmed the presence of MutL (MW_monomer_ ∼ 59 kDa).

### Calibration

#### Benchmark DNA samples

DNA oligomers with sequences from the benchmark single-molecule FRET study (51) were purchased from Integrated DNA Technologies (IDT) with an internal amino modifier C6 deoxythymidine (dT) at the sites for fluorescent labeling (see Supplementary Table S1 for sequences and Figure 1C for schematic). For labeling, we incubated each donor (D) or acceptor (A) oligomer with a 20x molar excess of the appropriate Atto-NHS ester dye (Sigma-Aldrich 92835 and 18373) in a fresh sodium bicarbonate buffer in 18.2 MΩ double-deionized water, ddiH_2_O (adjusted to pH 8.5 with 6M HCl) and reacted overnight at 4°C. The oligomers were purified with a ZYMO DNA oligo purification kit (D4060), and the UV/Vis absorbance spectrum was measured to determine labeling efficiency (See Supplementary Table S2). The labeled D and A oligomers were then annealed in equimolar amounts in 1x phosphate-buffered saline (PBS) by heating to 92°C for 4 minutes in a thermal cycler and cooling to room temperature at - 1°C per minute.

#### RNA-DNA hybrid substrates

RNA-DNA hybrid substrates designed for a nuclease cleavage assay were purchased from IDT fluorescently labeled with acceptor (Atto647N) and donor (Atto532 or AlexaFluor546 – AF546). The hybrid strand contained an acceptor dye on the 5’ RNA end (5’-/5ATTO647NN/rArGrA rArArU rArArG rGTA TCG GCA CGC TCG AGA TGA GGT ATT TCA CAC CTT AAG CCA GCC CCG ACA -3’), and the complementary DNA strand (5’-/5Biosg/TGT CGG GGC TGG CTT AAG GTG TGA AAT ACC TCA TCT CGA GCG TGC CGA TA/3ATTO550N/ -3’) contained a donor dye on the 3’ end and a biotin on the 5’ end. The donor-acceptor separation for this construct is 10 bases. Upon annealing the two oligomers by heating to 92°C for 4 minutes in a thermal cycler and cooling to room temperature at - 1°C per minute, we obtained a high FRET signal for the annealed substrate. We also purchased a variation of the oligomers that yielded medium FRET (5’-/5ATTO647NN/ rArGrA rArArA rArGrA rArArU rArArG AGA ATA TCG GCA CGC TCG TGA GGT ATT TCA CAC CTT AAG CCA GCC -3’ and 5’-/5Biosg/GGC TGG CTT AAG GTG TGA AAT ACC TCA CGA GCG TGC CGA TAT /iAlex546N/TCT -3’). The donor-acceptor separation for the second construct is 15 bases. Variations of the high and medium FRET substrates replaced uridine (Sub U) with adenine (Sub A) in the RNA-DNA hybrid strand.

#### Fluorescence correlation spectroscopy (FCS)

Individual detected photons are spectrally separated between the two SPAD detectors into green/donor (Channel 2) and red/acceptor (Channel 1) emission by dichroic and bandpass optical filters. Fluorescence intensity traces for each channel are generated by grouping sequential photon macrotimes into user-defined equal time bins (usually 1 millisecond). The fluorescence intensity traces for each channel can be autocorrelated (FCS), or the two signals can be cross-correlated (FCCS). The result is a correlation decay that is fit to an appropriate diffusion model to yield experimental parameters that characterize the diffusion characteristics of the fluorescently labeled species. We measured the diffusion characteristics for several approximately spherical particles. Fluorescent liposomes were prepared as described above, fluorescent beads were purchased from ThermoFisher (F8801 and F8784) and Bangs Laboratories Inc. (FSFR003), and proteins were expressed and fluorescently labeled as described above. QDs functionalized with streptavidin were purchased from Sigma-Aldrich (OCNQSS580). All samples were diluted to 10 nM in ddiH_2_O, a 20 μL droplet was pipetted onto a no.1 coverslip, and the laser was focused 40 μm into the droplet. For all measurements, the laser power was ∼ 6.5 kW/cm^2^ and measurements were acquired for 5 minutes.

For fitting to the Stokes-Einstein relation (equation 7), the radius of the liposomes was taken to be half the diameter of pores in the membranes used for extrusion. The radius of fluorescent beads and QDs was provided by the manufacturer. A cryo-EM reconstruction was used to determine the radius of Nsp15 (62). The Stokes radius of a MutL homolog was determined previously by sucrose density gradient centrifugation and gel filtration (63).

#### Pulsed interleaved excitation (PIE)-FRET

Solutions of annealed D-A labeled DNA were diluted to 10 nM in PBS. A 20 μL droplet was pipetted onto a no.1 coverslip, and the laser was focused 40 μm into the droplet. The power for each laser (531 and 636 nm) was adjusted until they were approximately equal in PIE mode. The total laser power density (red + green) was ∼ 3 kW/cm^2^ at 20 MHz repetition rate to ensure the full excited state decay was collected. For each sample, at least three measurements of 3 minutes each were collected, sampling thousands of molecules over the course of the acquisition.

Samples of individual and annealed RNA-DNA hybrid substrates were diluted to 5nM in a cleavage buffer (30 mM HEPES, 100 mM NaCl, 5 mM DTT, 5 mM MnCl_2_). A 40 μL droplet was pipetted onto a no.1 coverslip and the laser was focused 40 μm into the droplet. The power for each laser (531 and 636 nm) was adjusted until they were approximately equal in PIE mode. The laser power density (red + green) was ∼ 12 kW/cm^2^ at a 20 MHz repetition rate. Several 2-minute measurements were taken for all samples.

#### Preparation of functionalized microfluidic channels for single molecule surface attachment

The benchmark oligomers were purchased with a 5’ biotin modification on the acceptor strand (see Supplementary Table S1) for attachment to a homebuilt microfluidic chamber for imaging and spectroscopy. Quartz slides were predrilled prior to functionalization using 0.75 mm diamond-tipped drill bits (Triple Ripple) and a micro drill press. Slides and coverslips were cleaned by sonication in acetone, ethanol, and aqueous 1 M potassium hydroxide for 15 minutes each. Cleaned slides were stored in ddiH_2_O. To prepare the mixture of methoxy and biotinylated PEG-silane (mPEG and bPEG, respectively) for slide and coverslip functionalization, ∼ 20 mg of mPEG (Laysan Bio MPEG-SIL-2000, mean M_w_ of 2 kDa) was dissolved in 80 µL of ddiH_2_O and ∼ 2 mg of bPEG (Laysan Bio Biotin-PEG-SIL-3400, mean M_w_ of 3.4 kDa) was dissolved in 10 µL of ddiH_2_O. After mixing 1 µL of the bPEG solution with the mPEG solution, 40 µL of bPEG/mPEG mixture was sandwiched between a dried slide and coverslip. The assembly was reacted overnight in a dark, humid environment. The slides and coverslips were separated, rinsed with ddiH_2_O, and dried with pressurized air.

Once dried, another 40 µL of mPEG solution, prepared as above, was sandwiched between the slide and coverslip and allowed to react in a dark, humid environment at RT for ∼30 minutes. The slides and coverslips were then rinsed and dried once more before assembling them into homebuilt microfluidic chambers for singlemolecule measurements.

The microfluidic chambers were assembled by placing thin strips of double-sided tape between pairs of drilled holes on the slides to form individual channels of ∼ 20 µL volume. The coverslip was then placed on the tape so that the mPEG/bPEG functionalized sides of the coverslip and slide faced one another. The cover slip was carefully pressed down to ensure the tape adhered fully to both sides. The edges of the chambers were sealed by applying epoxy and allowed to set for 1 hour to overnight. Individual channels were washed three times with 100 μL of PBS. Next, 30 µL of 0.1 mg/mL streptavidin was injected into the chamber and allowed to bind to the biotinylated surface for 20 minutes. The chamber was washed again with PBS. Labeled, annealed biotinylated oligomers were added to the chamber at 10 pM and allowed to bind for 20 minutes. The chamber was rinsed three times to remove any unbound sample with 100 μL of PBS with added 2% glucose and 0.02% cyclooctatetraene for imaging. The addition of glucose is for a glucose oxidase/catalase oxygen scavenging system, which was not used in this study.

## RESULTS AND DISCUSSION

In confocal microscopy, fluorescence from the sample is excited with a focused laser beam and collected through a high numerical aperture objective lens. The signal passes through a dichroic mirror, and then a confocal pinhole for spatial filtering. The filtered fluorescence photons are directed to detectors with single photon sensitivity using optical elements. Spectral or polarization filters placed in the optical path divide the signal between multiple detectors. In our time-resolved fluorescence microscope with TCSPC, the detectors are connected to a fast event timer with picosecond resolution that tags each detected photon with a time relative to the start of the experiment (macrotime) and a time relative to the laser pulse rate (nanotime). TCSPC enables simultaneous detection of diffusion, fluorescence lifetime, and smFRET signals. The following sections present methods to validate FCS and PIE-FRET measurements. Obtaining accurate diffusion coefficients for biological molecules with FCS first requires careful characterization of the confocal volume. We determined the volume, *V*_*eff*_, and the aspect ratio, *κ*, of the confocal laser spot by using bead scanning and by measuring dyes with known diffusion coefficients (64). We then produced a calibration curve for the FCS measurements using approximately spherical nanoscale particles. Next, PIE-FRET histograms of diffusing DNA samples are presented, and we compare extracted FRET efficiencies for each sample to the previous benchmark study. Lastly, we compare the FRET efficiency values obtained for diffusing vs. surface attached benchmark DNA samples using intensity and lifetime-based analysis.

### Fluorescence Correlation Spectroscopy Calibration

Fluorescence correlation spectroscopy (FCS) is an established technique (11), but when combined with simultaneous PIE-FRET and fluorescence lifetime data, one gains extraordinary access to dynamic information across multiple timescales. In FCS, fluorescent molecules diffusing through a confocal volume cause fluctuations in the detected fluorescence intensity over time. One can imagine that fast-diffusing molecules will cause rapid changes in fluorescence intensity, while slow-diffusing molecules lead to more gradual changes.

These signal fluctuations can be characterized by performing an autocorrelation, *G*(*τ*),on the fluorescence intensity signal. Such a correlation calculates the similarity between the signal and a time-lagged version of itself by changing the lag time, *τ*, in the operation. With increasing *τ*, the time-lagged version becomes less similar to the original signal. Figure 2A shows a typical intensity trace for fluorescent liposomes diffusing through the confocal volume. An autocorrelation is performed on the data using equation 1 (65):

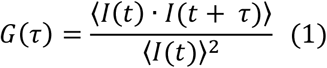

where *I*(*t*) is the fluorescence intensity over the acquisition time, *τ* is a lag time, and the brackets indicate an average over all time values, *t*. We obtain an autocorrelation curve that decays as the lag time increases (Figure 2B). Fitting the decay to an appropriate model yields experimental parameters that characterize the physical properties of the fluorescently labeled species (13). Here, we describe three diffusion models: pure diffusion, conformational, and triplet state. The use of each model depends on the sample properties. For all models, at *G*(0) the operation gives a maximum correlation amplitude as the signals (original and time-lagged) are identical. Fitting the decay to a diffusion model (12), *G*(0) is inversely related to the average number of molecules in the confocal volume at any instant in time, *N*. The lag time at which the autocorrelation amplitude decays to half its initial value is taken as the characteristic diffusion time, *τ*_*D*_, of the fluorescent species. Typical diffusion times for biomolecules are on the millisecond timescale. Equation 2 (65) is a pure diffusion model that can describe the motion of small fluorescent particles:

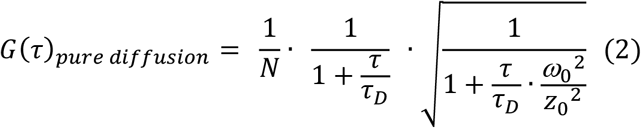

where *N* is the number of molecules in the confocal volume at any instant, *τ*_*D*_ is the characteristic diffusion time of the species, *ω*_0_ is the diameter of the laser beam waist, and *z*_0_ is the axial length of the laser beam (Figure 2C). The ratio 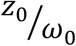 is the confocal volume aspect ratio, represented by *κ*. The aspect ratio can be measured directly or estimated, as described later. Once the *κ* value is determined, we can calculate the effective confocal volume as 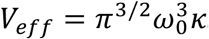.

**Figure 2:**
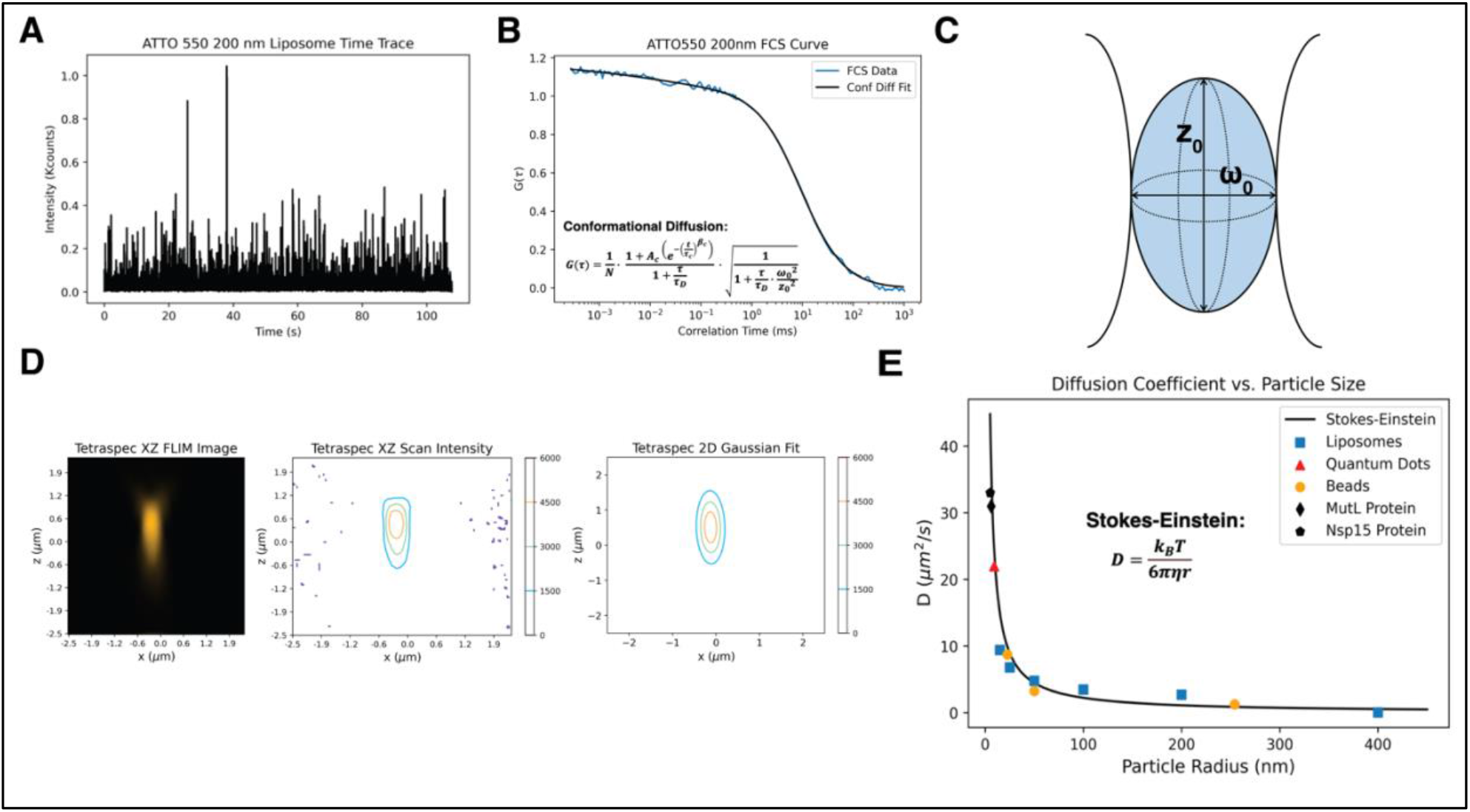
FCS calibration. **A)** The uncorrelated intensity data collected over time for Atto550 encapsulated in liposomes extruded with Whatman Nucleopore Track-Etch 200 nm membranes. **B)** The autocorrelated intensity data G(τ) (blue trace) and the conformational diffusion fit (black line, equation 3) for the 200 nm liposomes with encapsulated Atto550 dye. The fit parameters are: A_c_ = 0.136,τ_C_ = 0.032 μs,β_C_ = 0.302,τ_D_ = 9.2 ms,k = 6.49,N = 0.96, and D = 3.5μm^2^/s. **C)** Schematic of laser confocal volume in the XZ plane. ω_0_ is the diameter of the laser beam waist, and z_0_ is the axial length of the laser beam. The ratio 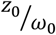 is the confocal volume aspect ratio, k. **D)** Bead image taken by scanning in the lateral (x) and axial (z) directions (left), a contour plot of the image intensity (center), and the 2D elliptical Gaussian fit (right, equation 6). **E)** Diffusion coefficients for several approximately spherical particles of different radii were determined using FCS with our calibration parameters. The Stokes-Einstein diffusion model (black line, equation 7) was fit to the data. We used η = 0.0009608 Pa ∗ s, and T = 294 K in the Stokes-Einstein model. χ^2^ = 10.49 for the fit to the experimental data.

For samples other than small organic dye molecules, a physical model besides the pure diffusion model (equation 2) is often required to adequately fit the experimental data. To account for conformational changes that occur in larger biomolecules (i.e., proteins, nucleic acids, and lipids) while they are in the confocal volume, a stretched exponential component is added to the pure diffusion equation to model transitions between two conformation states (equation 3) (65):

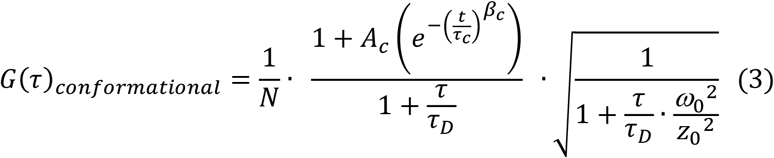

where parameters previously defined for equation 2 are the same, *A*_*C*_ is a pre-exponential factor, *τ*_*C*_ is a relaxation time for reversible transitions between the two states, and *β*_*c*_ is a stretch parameter.

Some fluorescent molecules enter non-emissive states called triplet states (66). Such photophysical behavior is typical for red-emitting dyes, and these excited states have longer fluorescence lifetimes than singlet state photon emission. The behavior of these dark states in molecules can be accounted for in autocorrelation curves by separating the average proportion of the molecules that are excited into the triplet state and scaling the triplet population by the characteristic time, *τ*_*T*_, that a molecule resides in the triplet state. The triplet state model is shown in equation 4 (65):

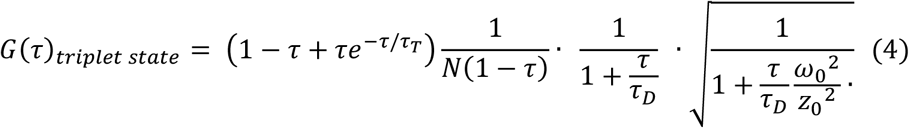

To convert the characteristic diffusion time, *τ*_*D*_, extracted from the three diffusion models above to a diffusion coefficient, *D*, in area per unit time, we can use equation 5 (65):

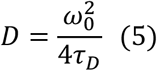

where *ω*_0_ is the diameter of the laser beam waist. We can compare this converted *D* to the theoretical diffusion of spherical particles governed by the Stokes-Einstein equation as outlined later. In addition, *D* has a larger dynamic range for identifying biomolecules based on differences in size. The accurate determination of *D* from equation 5 depends on a rigorous characterization of the three-dimensional confocal volume, which we have approached in two ways including bead imaging and fitting FCS curves of organic dyes (64). In the first method, we scanned fixed fluorescent beads in perpendicular planes. These beads are smaller than the image resolution and act as point sources of light. The resulting scan intensity profiles in each plane: *XY, XZ*, and *YZ*, are fit to a 2D elliptical Gaussian to obtain an intensity profile using equation 6:

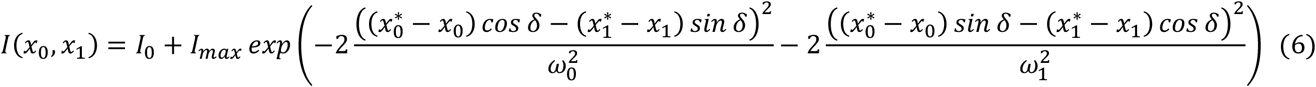

where 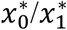,and *δ* describe the center and angle of the 2D elliptical Gaussian, respectively; and ω_0_/ω_1_ describe the widths of the 2D elliptical Gaussian. We imaged a prepared slide (PicoQuant) containing 100 nm Tetraspek® beads on a coverslip and fit the intensity profiles to determine *V*_*eff*_, *ω*_0_, *z*_0_ and *κ*. Figure 2C shows the schematic of a focused laser beam as a side profile. As discussed previously, *k* is 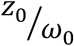,or aspect ratio. An example image of a single bead taken in the *XZ* plane is shown in Figure 2D (left). A contour plot of the image is in Figure 2D (center), and the fit to equation 6 is in Figure 2D (right). The calculated values of *V*_*eff*_, *ω*_0_, and *κ* for our three laser wavelengths are in Supplementary Table S3 (top).

The second method of characterizing the confocal volume relies on fitting measured FCS curves of organic dyes with known diffusion coefficients to a model function. We measured the FCS signal of each sample solution by pipetting a droplet onto a thin glass coverslip, focusing into the droplet, stopping the beam at a single point, and collecting fluorescence photons for several minutes. During the acquisition, thousands of molecules diffuse through the confocal volume, each emitting photon bursts upon successive excitations by the pulsed laser source. The fluorescence intensity traces obtained were autocorrelated and for consistency, the triplet state model (equation 4) was used for all fits to extract *τ*_*D*_,*τ*_*T*_ and *κ* parameters. The fit parameters were used to calculate the effective volume by substituting equation 5 into the effective volume equation to obtain:

*V*_*eff*_ = *π*^3/2^(4*Dτ*_*D*_)^3/2^*κ*. Supplementary Table S3 (bottom) displays the resulting aspect ratio and calculated effective volumes from the FCS triplet state fits for each laser wavelength. Organic dyes used for each wavelength are indicated in the table. Comparing the two approaches, the calculated effective volumes for each wavelength are similar, while the aspect ratio measured from FCS curve fitting is roughly twice that directly measured from bead scanning. Fitting experimental data with either pair of calibration values for the selected laser and approach often yields similar results, but using the *κ* and *V*_*eff*_ values obtained from FCS measurements gives more consistent results for the various diffusing species we measure. The bead scanning method, however, is a more accurate measure of the true confocal laser spot, or point spread function, which would be useful for deconvolving an image.

The diffusion models described above (equations 2,3,4) are used to extract physical and photophysical characteristics of the diffusing species by fitting the experimental data. Theoretically, the diffusion of small spherical particles in a solution can be described by the Stokes-Einstein relation (57), equation 7:

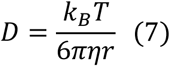

where *D* is the diffusion coefficient, *k*_*B*_ is the Boltzmann constant, *T* is temperature, *η* is the viscosity, and *r* is the particle radius. Based on this relationship, a smaller particle will have a larger diffusion coefficient, and vice versa. To test for convergence of theory and experiment, we prepared solutions of approximately spherical particles including fluorescent liposomes, quantum dots, and beads. We also determined diffusion coefficients for two labeled proteins. Nsp15 and *Taq* MutL proteins were expressed, purified, and fluorescently labeled (see Materials and Methods section). FCS curves were generated and fit to a suitable model (either pure diffusion or conformational as described above) to extract a diffusion coefficient, *D*, using equation 5. We plotted the diffusion coefficient versus the particle radius and fit the data to the Stokes-Einstein equation (Figure 2E).

Fitting the experimental data to the Stokes-Einstein curve yields a great fit (*χ*^2^ = 10.49), indicating that the theoretical model accurately describes the diffusion of the approximately spherical particles. Our rigorous analysis is therefore capable of distinguishing biomolecules that vary in size. This approach can be applied to determining oligomeric states of proteins, or cleavage of a nucleic acid substrate by a nuclease, for example.

#### FCS characterization of benchmark DNA samples

Prior to calibration of the PIE-FRET signal with the benchmark DNA samples, we measured the physical and photophysical properties of the samples with FCS. Each benchmark DNA oligomer is 38 bases with amino dT sites for labeling, and the acceptor strand contains a 5’ biotin (see Supplementary Table S1 for sequences). The determined FCS calibration values were applied to calculate the diffusion coefficient, *D*, of the fluorescently labeled benchmark DNA samples (Figure 1C), measuring *D* at each step of the preparation process. First, the diffusion coefficients for the Atto550-NHS ester and Atto647N-NHS ester dyes used for labeling were calculated, obtaining average values of 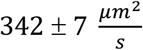 and 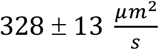, respectively. The diffusion coefficient for the Atto647N-labeled acceptor DNA strand, which also contains a biotin molecule, was 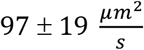. This significant decrease in *D* compared to the free dye molecule is due to the large size of the covalently attached DNA molecule, which causes the dye to diffuse more slowly. For the lo, mid, and hi Atto550-labeled donor strands, *D* ranges from an average of 130 to 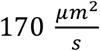, consistent with a slightly smaller construct as it does not contain a biotin. The three annealed donor-acceptor (D-A) duplex DNA substrates had diffusion coefficients ranging from an average of 85 to 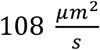. This variation in the measurement may be due to incomplete annealing. Representative experimental autocorrelation decays with fits for each sample described above are presented in Supplementary Figure S1, and the average values are summarized in Supplementary Table S4. Although the DNA molecules deviate from spherical particles, the trends and consistency of the data indicate that FCS, when carefully calibrated, can reliably distinguish between site-specifically labeled linear DNA molecules of different sizes.

### Pulsed Interleaved Excitation (PIE)-FRET Calibration

FRET is a photophysical effect that results from a fluorescent donor (D) molecule in the excited state transferring energy via dipole-dipole coupling to a nearby acceptor (A) molecule with favorable spectral characteristics (67,68). The dye spectral properties and optical setup are selected such that one can excite a donor fluorophore without directly exciting the acceptor, thus any acceptor emission is from resonant energy transfer. The energy transfer efficiency depends on the sixth power of distance *R* between donor and acceptor as *E*_*FRET*_ = [1 + (*R*/*R*_0_)^6^]^−1^, where *R*_0_ is the Förster radius at which *E*_*FRET*_ = 50% for a specific donor-acceptor (D-A) pair. This strong distance sensitivity makes FRET a nanoscale ruler that can detect distances between 3 to 10 nanometers (68). The donor and acceptor molecules can be covalently attached to biomolecules to report on the proximity between two sites.

Accurate recovery of the true FRET efficiency from experimental donor and acceptor intensity data relies on the determination of three correction factors: *α, δ*, and *γ* (46). The correction factor *α* accounts for spectral cross-talk from donor fluorescence leakage into the acceptor channel while the correction factor *δ* accounts for the direct excitation of the acceptor with the donor laser. The *γ* correction factor accounts for relative differences in the quantum yields and detection efficiencies of the donor and acceptor dyes (46,51,69). A previous smFRET benchmark study (51) outlined a robust correction procedure for all three factors and focused on intensity-based measurements where FRET efficiency was calculated from donor and acceptor photon counts after donor excitation. Here, we focus on the use of pulsed interleaved excitation (PIE or nsALEX) (47-50) to sequentially excite donor and acceptor molecules directly as they diffuse through the confocal volume. The quasi-simultaneous detection of donor, acceptor, and FRET signals with ALEX/PIE removes analysis uncertainty by confirming the presence of an active acceptor, and ensuring a single donor-acceptor pair is probed. Using a microscope system powered by TCSPC coupled with PIE, we can simultaneously monitor diffusion characteristics, structural dynamics, and interactions of single biomolecules diffusing through the confocal observation volume.

The residence time of the molecules in the confocal volume is on the millisecond timescale as discussed in the FCS section, while the time delay between sequential picosecond laser pulses is on the nanosecond timescale (Figure 1A). Thus, individual molecules undergo thousands of cycles of excitation and emission while in the confocal volume, and the macrotimes and nanotimes are binned to obtain fluorescence intensity traces and excited state decays for the sub-ensemble. The association of each burst of photons with the excitation pulse and emission channel leads to quasi-simultaneous monitoring of the intensity and lifetime of donor and acceptor molecules while in the confocal volume. In addition, the PIE method can be combined with multiparameter fluorescence detection (MFD) (49) to obtain fluorescence anisotropy, but that is not explored in this study. PIE offers several possibilities to accurately determine correction factors using intensity and lifetime-based measurements to ultimately uncover the true FRET efficiency.

Using the PIE configuration on our microscope system, the donor and acceptor dyes are excited by rapidly switching (or interleaving) the 531 and 636 nm laser pulses on the nanosecond time scale (47) (Figure 3A). The donor and acceptor emission are separated by a 635 nm long pass dichroic filter placed between the two detectors and bandpass filters in front of each detector isolate acceptor (690/70, Channel 1) and donor (582/64, Channel 2) emission. The upper left quadrant in Figure 3A illustrates the 636 nm laser pulse excitation (*ex*) of the acceptor molecule (*A*_*ex*,_ Time gate 1). The acceptor emission (*em*) is subsequently detected in Channel 1 (*A*_*ex*_*A*_*em*_), and the result of multiple cycles of excitation and emission is indicated by a filled decay curve. The decay is generated by binning the nanotimes for all detected photons. The lower left quadrant is empty because *A*_*ex*_*A*_*em*_ is red-shifted and should not be detected in Channel 2 which senses shorter-wavelength donor emission. At a repetition rate of 40 MHz in PIE mode, the 531 nm laser pulse occurs 25 nanoseconds later, exciting the donor molecule (*D*_*ex*,_ Time gate 2). The upper right quadrant illustrates the laser pulse and the signal detected in Channel 1 after many cycles of *D*_*ex*_, which consists of FRET if there is a nearby acceptor (*D*_*ex*_*A*_*em*_) and some donor leakage. The lower right quadrant depicts donor emission detected in Channel 2 after many cycles of donor excitation (*D*_*ex*_*D*_*em*_). The intensity (and fluorescence lifetime) of the donor emission is reduced due to FRET. As mentioned previously, FRET efficiency can be calculated with intensity or fluorescence lifetime-based measurements. While we have used excited state decays to illustrate PIE, we determined the *α* and *δ* correction factors with fluorescence lifetime and intensity data, which we compare below.

**Figure 3:**
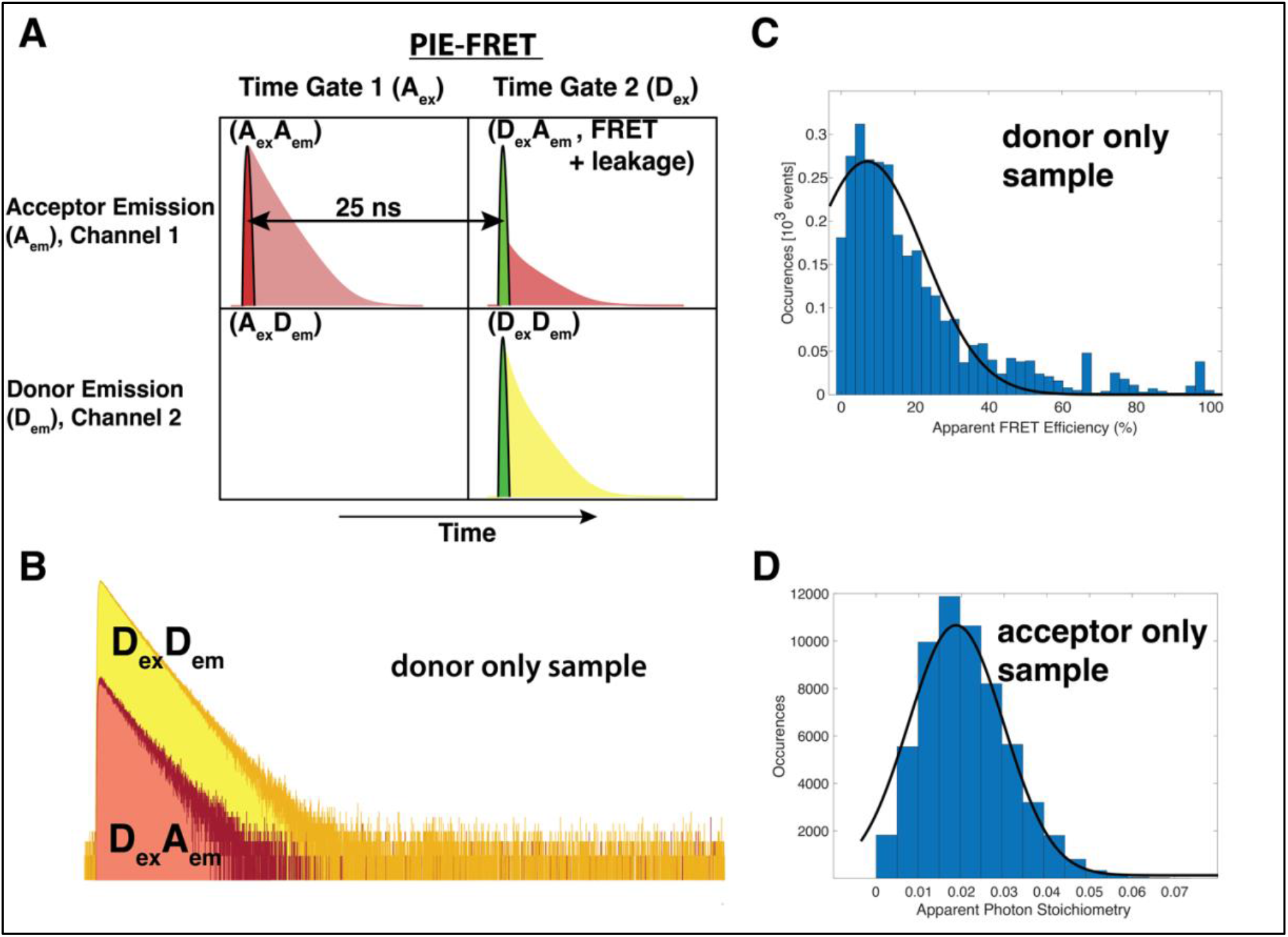
PIE-FRET schematics. **A)** Principle of pulsed interleaved excitation – PIE. The red laser pulses in the first time gate, and photons from acceptor molecules are detected only in Channel 1. The excited state decay results from creating a histogram of photon arrival nanotimes. After a 25 ns delay and complete decay of the acceptor excited state, the green laser pulses to excite the donor molecules in the second time gate. Donor emission is detected in Channel 2, and if FRET occurs acceptor emission is detected in Channel 1 during the second time gate (in addition to some spectral crosstalk or leakage that we account for by determining the correction factor α). **B)** Example data used to calculate the donor leakage correction factor (α) of a donor only mid-Atto550 sample (Supplementary Table S1) with the photon count method (equation 8) applied to fluorescence lifetime data. **C)** Representative histogram (blue bars) obtained from calculating the average apparent fluorescence intensity-based FRET efficiency, (⟨E_app_⟩), using equation 9 for a donor only hi-Atto550 sample to calculate α with equation 10 after fitting the histogram to a Gaussian function of the form 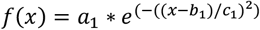 (black line). The fit parameters are: a1 = 0.26, b1 = 5.5 (%), c1 = 23 (%) (R^2^ = 0.93) **D)** Representative histogram (blue bars) obtained from calculating the average fluorescence intensity-based apparent photon stoichiometry, (⟨S_app_⟩), of an acceptor only sample using equation 12. The direct excitation factor δ was calculated with equation 13 after fitting the histogram to a Gaussian function. The fit parameters are: a1 = 1.2E4, b1 = 0.017, c1 = 0.015 (R^2^ = 0.99).

We estimated the *α* correction factor with two different approaches using photon counts or Gaussian curve fitting. Atto550 free dye and Atto550-NHS ester labeled benchmark oligomers were used as donor-only samples diffusing in a droplet solution. In the first method, we calculated the ratio of the photon counts (from nanotimes) detected in the acceptor Channel 1 to the photon counts detected in the donor Channel 2 for the donor-only measurement, effectively calculating the ratio of the area under each curve in Figure 3B to determine *α*. The data in Figure 3B result from a fluorescence lifetime measurement in PIE mode after donor excitation (*D*_*ex*_,Time gate 2, 531 nm) of the Atto550-donor mid strand. It is important that the excited state completely decays during the time gate, thus a suitable repetition rate should be selected. The *α* calculation from photon counts can be represented using equation 8:

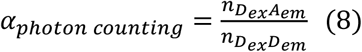

where 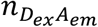 represents the photon counts detected in the acceptor channel (*A*_*em*_) during only the donor excitation time gate (*D*_*ex*_) and 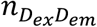 represents the photon counts detected in the donor channel (*D*_*em*_) during only the donor excitation time gate (*D*_*ex*_).

In a second method to determine *α* using an intensity-based measurement, the PIE-FRET signal was measured for molecules diffusing in solution by exciting the sample in PIE mode and collecting data for several minutes. As individual fluorophores diffuse through the confocal volume, they emit photon bursts. The macrotimes for photons detected in each channel were binned into equal 1-millisecond bins, and a burst threshold was determined using a burst search algorithm to identify individual diffusion events above the background signal (24). The background was subtracted to generate a fluorescence intensity (*I*) time trace for each channel. We performed the following apparent FRET efficiency 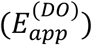 calculation on the fluorescence intensity signal from the donor-only (DO) species using equation 9 (51):

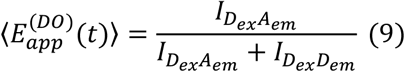

There is no acceptor present in the sample, thus actual FRET is not possible. A histogram of 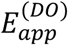 calculated over the entire fluorescence intensity trace was generated and fit to a Gaussian function of the form 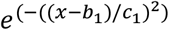 (Figure 3C). We took the Gaussian peak value (b_1_ = 5.5 %) as the average apparent FRET efficiency, 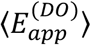, which was used in equation 10 (51) to calculate *α*:

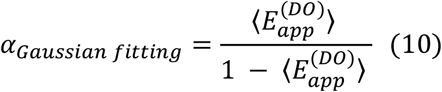

We achieved good agreement between the two methods, as shown in Supplementary Table S5, with fluorescence lifetime and intensity-based strategies yielding similar results for *α*. We note that the fluorescence lifetime measurement analysis is faster and more straightforward by comparison. The spectral cross talk is very similar for Atto550 free dye and the labeled benchmark DNA oligomers, suggesting a negligible impact of the DNA on the Atto550 spectral properties.

Similarly, we used fluorescence lifetime and intensity methods to calculate *δ*,the correction factor accounting for direct excitation of the acceptor with the donor laser. We estimated the *δ* correction factor by taking the ratio of the detected photon counts from *D*_*ex*_ *A*_*em*_ to the detected photon counts from *A*_*ex*_ *A*_*em*_ for an acceptor-only (AO) fluorescence lifetime measurement collected in PIE mode. The 531 and 636 nm laser power was adjusted such that they were equal. This calculation can be represented using equation 11:

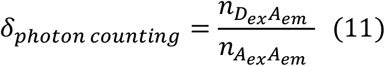

where 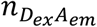 represents the photon counts detected in the acceptor channel (*A*_*em*_) during only the donor excitation time gate (*D*_*ex*_) and 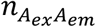 represents the photon counts detected in the acceptor channel (*A*_*em*_) during only the acceptor excitation time gate (*A*_*ex*_). The fluorescence decays are not shown, but the calculated values for *δ* are in Supplementary Table S5.

We also calculated *δ* by taking the acceptor-only species fluorescence intensity time trace with background corrections and calculating the average photon stoichiometry, *S*, over time using equation 12 (51):

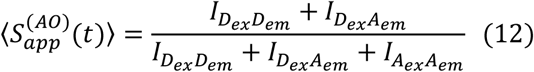

The photon stoichiometry is the ratio of photons detected in both donor and acceptor channels after donor excitation to the sum of all detected photons after donor and direct acceptor excitation in PIE mode. For the acceptor-only sample, there should not be a *D*_*ex*_ *D*_*em*_ signal. With a D-A sample that undergoes FRET, the stoichiometry ratio enables one to distinguish between subpopulations of donor-only, acceptor-only, and D-A pairs as described in detail later. A histogram of 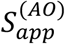 calculated over the entire fluorescence intensity trace was generated and fit to a Gaussian function (Figure 3D). We took the Gaussian peak value, b_1_ = 0.017, as the average apparent photon stoichiometry 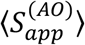, which was used in equation 13 (51) to calculate *δ*:

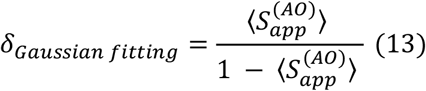

Supplementary Table S5 includes the average *δ* values for each sample measured which show excellent agreement between the two methods. We also determined the *α* and *δ* values for an RNA-DNA hybrid strand labeled with Atto647N, and its complementary DNA strand labeled with AlexaFluor546. The values are presented in Supplementary Table S5 as well (*AF546-DNA complement to RNA/DNA hybrid* and *Atto647N-RNA/DNA hybrid*, see Materials and Methods for sequences). The *α* and *δ* correction factors were then used to calibrate the PIE-FRET signals from the annealed D-A benchmark DNA duplexes and RNA/DNA hybrids.

To calibrate the PIE-FRET signal, we measured the three annealed D-A benchmark DNA samples (lo, mid, and hi FRET), simultaneously acquiring FCS, fluorescence lifetime, and PIE-FRET data. Representative fluorescence lifetime and diffusion coefficient data obtained for free dyes, single-stranded, and annealed DNA samples are presented in Supplementary Figure S1 and Supplementary Table S4 as described previously. PIE enables precise determination of the FRET efficiency by removing the contribution of complexes that do not contain an active acceptor molecule. Using the separated *D*_*ex*_ *D*_*em*_,*D*_*ex*_ *A*_*em*_ (*FRET*) and *A*_*ex*_ *A*_*em*_ signals from PIE, we can calculate the corrected PIE-FRET efficiency, *E*, with equation 14 (51):

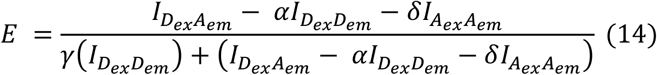

and the photon stoichiometry, *S*, using equation 15 (51):

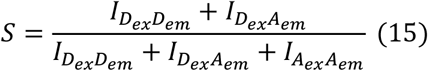

where 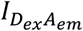 is the background-corrected intensity for the acceptor due to FRET, 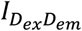 is due to direct donor excitation (531 nm), and 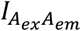 is due to direct acceptor excitation (636 nm). Parameters *α, δ*, and *γ* are correction factors introduced earlier. For acceptor-only labeled molecules *S* = 0, for donor-only labeled molecules *S* = 1, and for donor-acceptor pairs *S* = 0.5. In practice, 0.3 < *S* < 0.8 can be attributed to donor-acceptor labeled species (45). Plotting the PIE-FRET efficiency *E* versus the photon stoichiometry *S* gives 2D *E-S* histograms where subpopulations can be visualized.

Two-dimensional *E-S* histograms for benchmark DNA were produced using the open-source PIE Analysis with MATLAB (PAM) software package, where a dual channel burst search (DCBS) was used for burst selection. PIE-FRET analysis eliminates donor-only and acceptor-only artifacts such that analysis is focused on the subpopulation of labeled substrates that contain one donor and one acceptor (0.3 ≤ *S* ≤ 0.8). We plotted two-dimensional *E-S* histograms for each sample before correction (Figure 4 A, C, E). In each *E-S* histogram (45), the x-axis is intensity-based PIE-FRET efficiency, *E*, calculated using equation 14, the y-axis is photon stoichiometry, *S*, calculated using equation 15, and occurrence histograms for stoichiometry (gray horizontal bars) and PIE-FRET (gray vertical bars) are shown on each axis.

**Figure 4:**
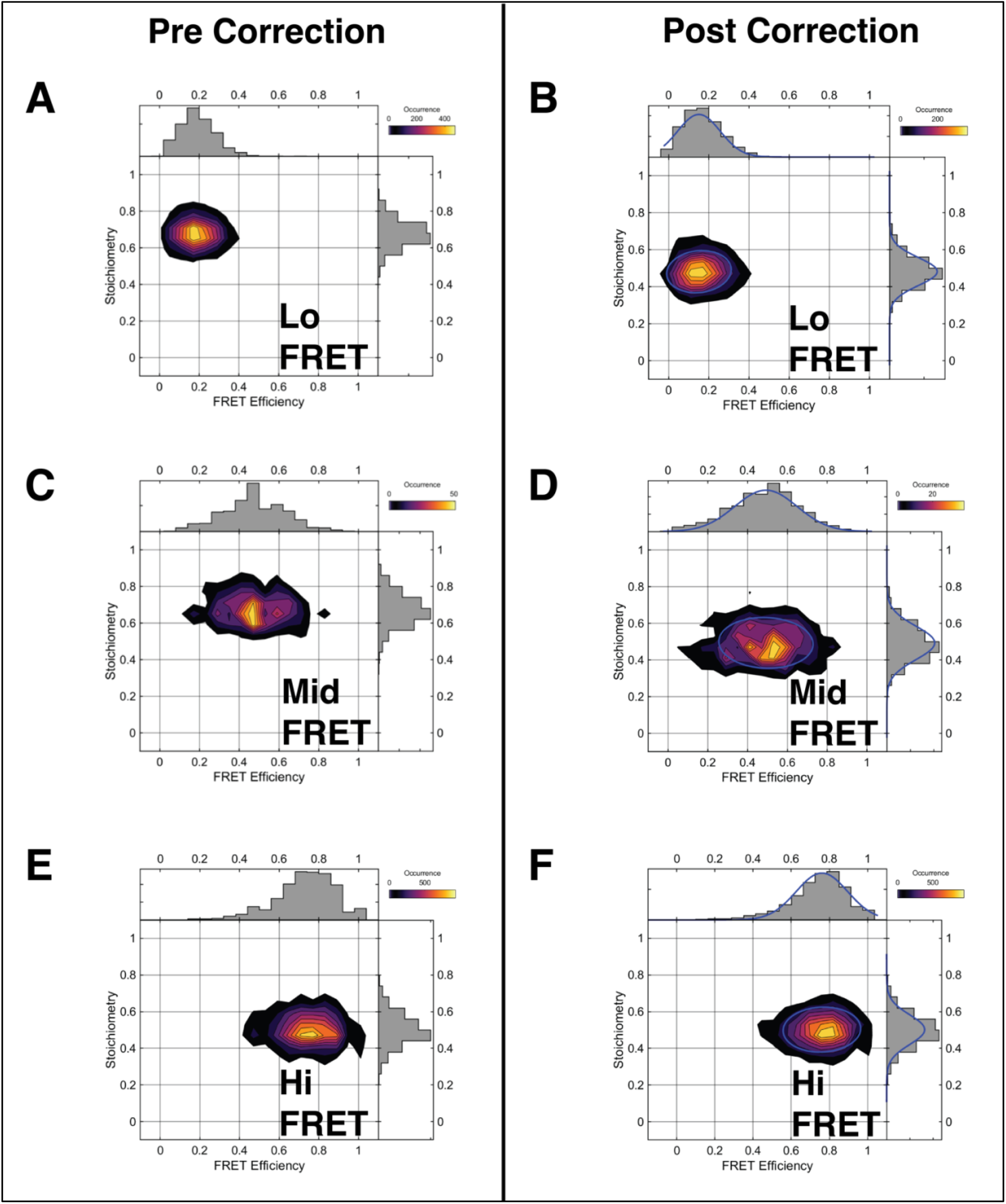
PIE-FRET calibration using lo, mid, and hi FRET benchmark DNA samples that were fluorescently labeled and annealed. All samples were diluted to 10 nM in PBS (20 nM for the hi FRET sample) and three 3-minute acquisitions were aggregated for analysis in the PAM software. In each 2D E-S histogram, the x-axis is PIE-FRET efficiency (E, vertical bars) calculated with equation 14 and the y-axis is photon stoichiometry (S, horizontal bars) calculated using equation 15. **A), C), E)** Lo, mid, and hi FRET samples, respectively, precorrection for donor leakage (α), direct acceptor excitation (δ), gamma (γ), and beta (β). **B), D), F)** Lo, mid, and hi FRET samples, respectively, postcorrection for donor leakage (α), direct acceptor excitation (δ), gamma (γ), and beta (β). The blue curves are Gaussian fits, and the fit parameters are summarized in Supplementary Table S6.

To correct the *E-S* histograms, the averages of correction factors *α* = 0.06 and *δ* = 0.02 (Supplementary Table S5) determined with the DO and AO samples as described above were applied. We then used the PAM software to determine *γ* and *β* correction factors. There is some uncertainty in determining the gamma factor for diffusing confocal experiments (9,10). An average gamma value of *γ* = 0.88 was determined from the linear fit of a burst-wise plot of *1/S* vs. *E* (46). The *β* factor, which accounts for different excitation efficiencies of donor and acceptor fluorophores, was also determined with the *1/S* vs. *E* plot. The corrected *E-S* histograms are shown in Figure 4 B, D, F. The PIE-FRET efficiency histograms were fit to a Gaussian function, and the parameters are summarized in Supplementary Figure S6. We measured and analyzed the hi FRET sample two ways to compare the quality of the data: high concentration/short acquisition, and low concentration/long acquisition, and the results are presented in Supplementary Figure S2. In the next section, we calculate gamma with fluorescence trajectories from single surface-attached DNA duplexes.

Each PIE-FRET efficiency histogram (vertical bars below two-dimensional *E-S* histograms in Figure 4) was fit to a Gaussian function to extract average FRET values for each sample. The Gaussian fits and parameters are shown in Supplementary Figure S2 and Supplementary Table S6. The average FRET efficiency, ⟨*E*⟩, for each of our labeled DNA samples after *α* and *δ* correction is - lo FRET: 0.15; mid FRET: 0.49; and hi FRET: 0.76. The FRET values for the Atto550/Atto647N labeled DNA samples reported in the previous benchmark study were-lo FRET: 0.15 ± 0.02; mid FRET: 0.56 ± 0.03; and hi FRET: 0.76 ± 0.015 (51). For clarity, our oligomers have the same sequence as the substrates from the benchmark study, but we fluorescently labeled and annealed our DNA in house as described in the Materials and Methods section. A direct comparison of the FRET efficiency values for our samples and the findings from the benchmark study demonstrate excellent agreement. The PIE-FRET histograms are relatively broad perhaps due in part to quantum yield variations of the acceptor dye that depend on donor – acceptor distance (70) and the short acquisition time (see Supplementary Figure S2 for example data collected for a longer period of time). We provide direct evidence for such an influence in the next section as we observe a trend in the fluorescence lifetime of the *D*_*ex*_*A*_*em*_ (*FRET*) signal between the lo, mid, and hi FRET samples. The corresponding distance histograms for the PIE-FRET data in Figure 4 are in Supplementary Figure S3 with corresponding calculations for the average distance between dyes based on a literature value for *R*_0_. The corrected PIE-FRET signals for the annealed RNA/DNA hybrid substrates were also calculated using the correction factors in Supplementary Table S5. The representative 2D *E-S* histograms, 1D PIE-FRET efficiency histograms, and FCS curves are shown in Supplementary Figure S4. Compared to the benchmark oligomers that contain 38 bases, the two RNA/DNA hybrid oligomers are 60 bases, while the DNA complements are 45 and 50 bases (see Materials and Methods). We observe much smaller diffusion coefficients of 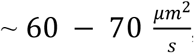, consistent with a larger nucleic acid molecule.

The fast laser scanning confocal configuration of our microscope also enables rapid imaging of single molecules attached to a surface, producing FLIM images that not only report the location of molecules, but also the fluorescence lifetime. Since FRET can be detected by calculating changes in the donor fluorescence lifetime, FLIM-FRET images can report on the energy transfer efficiency. Individual molecules in the images may be selected to probe single complexes over long periods of time and at high temporal resolution (until photobleaching occurs). We attached the annealed benchmark DNA samples to a surface and collected FLIM-FRET images and single-molecule fluorescence intensity traces, then compared the calculated FRET efficiency to the diffusing measurements.

### Single-Molecule FLIM-FRET Imaging

The benchmark oligomers were purchased with a 5’ biotin modification for surface attachment to homebuilt microfluidic channels. The channels were assembled as described in the Materials and Methods section, and the annealed biotinylated DNA duplexes were attached to the microscope slide and imaged with fluorescence lifetime imaging microscopy (FLIM) in PIE mode. We obtain *D*_*ex*_ *D*_*em*_, *D*_*ex*_ *A*_*em*_ (FRET), and *A*_*ex*_ *A*_*em*_ images with a single acquisition. Figure 5A shows the single molecule images for annealed lo, mid, and hi FRET samples. The color scale of the pixels in a FLIM-FRET image indicates the fluorescence lifetime, which changes for the donor when FRET occurs. Individual DNA duplexes indicated by a bright spot in the image were selected, and the fluorescence intensity time trace was collected at a single location.

**Figure 5:**
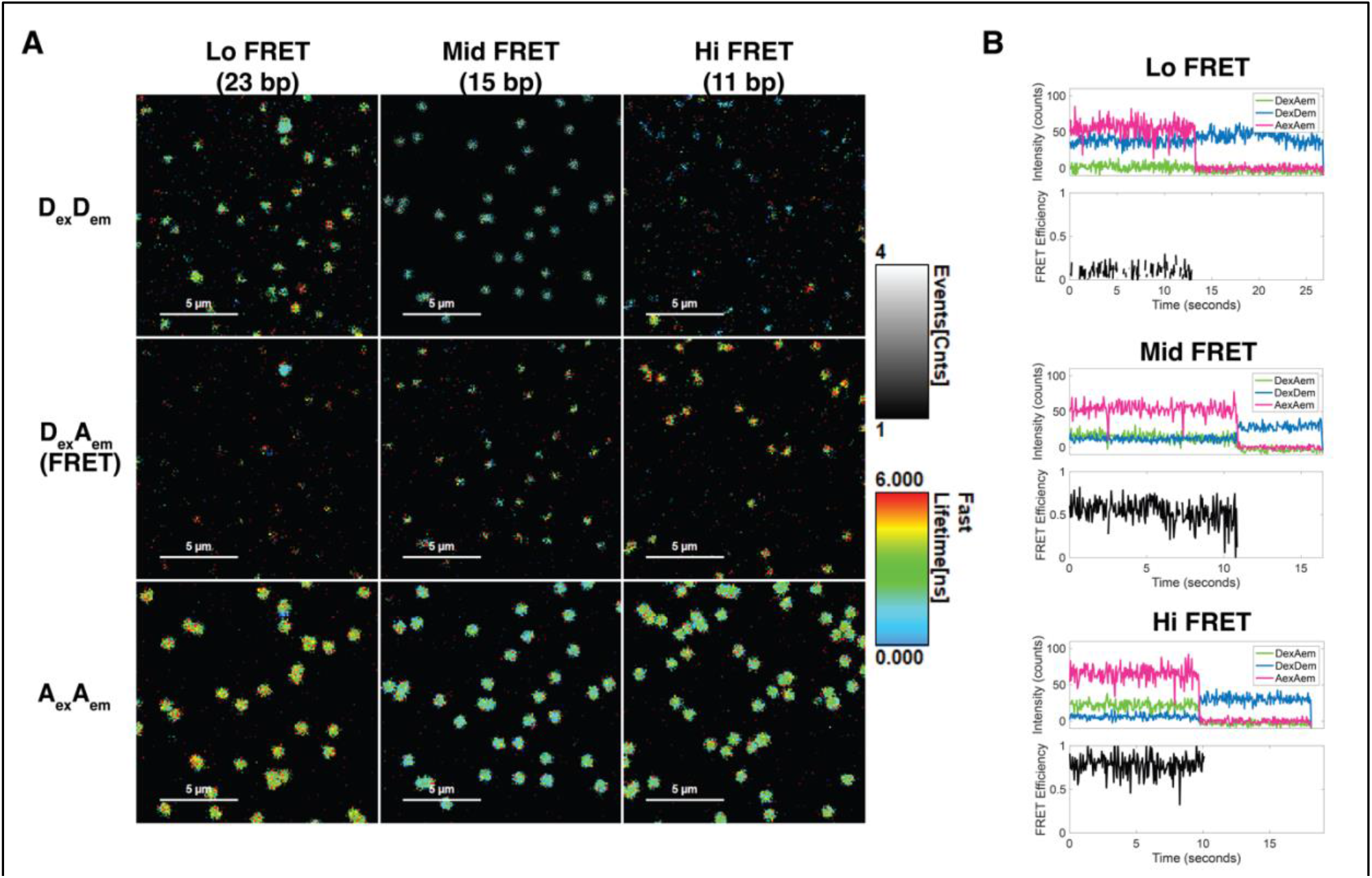
FLIM-FRET single molecule images and traces of benchmark substrates. **A)** D_ex_D_em_, D_ex_A_em_ (FRET), and A_ex_A_em_ images obtained simultaneously with PIE for each sample (lo, mid, and hi FRET). The brightness of each spot is related to the fluorescence intensity, indicated by the gray scale bar. Higher photon counts are brighter in the image, while lower photon counts are darker. The color of each pixel in the image indicates the Fast fluorescence lifetime at that pixel, which is the average arrival nanotime. Red represents a longer lifetime and blue is a shorter lifetime. **B)** Individual bright spots correspond to single fluorescently labeled annealed benchmark DNA molecules. Single-molecule fluorescence intensity traces for individual duplexes are obtained by clicking on a bright spot and collecting photons from that location. The photon macrotimes were then binned in 50 ms bins to generate the traces. In the traces for individual lo, mid, and hi FRET duplexes, D_ex_D_em_ is blue, D_ex_A_em_ (FRET) is green, and A_ex_A_em_ is magenta. The FRET efficiency (black line) below each set of traces was calculated with equation 14 using the correction factors in Supplementary Table S5. The photobleach step for the acceptor, which is followed by recovery of the donor fluorescence, is apparent in each trace. The γ correction factor was calculated with equation 16 considering the photobleach step.

Each bright spot in Figure 5A is a single DNA duplex (a few brighter spots may be 2-3 molecules). Due to the energy transfer that occurs in the FRET interaction between donor and acceptor dyes in an annealed substrate, we expect the donor intensity and fluorescence lifetime to be impacted by the distance from the acceptor dye. The brightness of each spot indicates the molecule fluorescence intensity, and the color of the pixels in each image indicate the fluorescence lifetime at that location, thus we can directly infer energy transfer from analyzing a set of FLIM-FRET images.

From left (lo FRET) to right (hi FRET) in Figure 5A, the donor signal (*D*_*ex*_ *D*_*em*_) decreases in intensity (represented by photon counts on a pixel-by-pixel basis). The gray scale bar indicates pixel brightness, with darker areas being less intense and brighter more intense. The fluorescence lifetime of the molecules is represented by the Fast Lifetime in nanoseconds (see color bar), which is the average nanotime of the photons on a on a pixel-by-pixel basis. From left (lo FRET) to right (hi FRET), the fluorescence lifetime of the donor molecules (*D*_*ex*_*D*_*em*_) also decreases (green color to blue color). As the donor and acceptor dye become closer (left to right), we also expect the acceptor signal due to FRET (*D*_*ex*_ *A*_*em*_) to increase as is observed in the middle set of images. Direct excitation of the acceptor dye in PIE mode (*A*_*ex*_ *A*_*em*_) varies slightly across the three samples (lower images).

After collecting a FLIM-FRET image, we clicked on individual bright spots and collected singlemolecule fluorescence intensity traces. A representative donor and acceptor intensity trace for each benchmark DNA sample is shown in Figure 5B. The FRET efficiency (black line) below each trace was calculated with equation 14 using the *α* and *δ* correction factors in Supplementary Table S5 determined from diffusing measurements. In each trace, the FRET signal vanishes after the acceptor photobleaches in one step. We calculated the gamma correction factor, *γ*, for surface attached molecules (69,71,72) with equation 16:

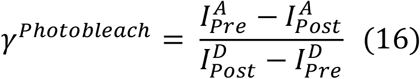

where 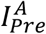 is the average intensity of the acceptor molecule prior to the photobleaching step, 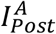 is the average acceptor intensity after the bleach, and 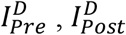 are the same values for the donor molecule. The *γ* factors for the lo, mid, and hi FRET traces (Figure 5B) were: 0.74, 0.59, and 0.64, respectively. The slight variation in the *γ* factor reflects the heterogeneity in single particles and their local environment, which may be obscured in diffusing measurements. For comparison, the average gamma factor determined with the diffusing PIE-FRET measurements was 0.88. The calculated FRET efficiencies from single-molecule fluorescence intensity traces using equation 14 for the lo, mid, and hi FRET traces were: 0.11, 0.54, and 0.68, respectively. There is strong agreement between values obtained from the diffusing PIE-FRET experiments conducted on the same samples (lo FRET: 0.14; mid FRET: 0.51; and hi FRET: 0.74). The fluorescence lifetime data can also be used to calculate FRET.

The photophysical process of relaxation from an excited state is influenced by radiative (photon-producing) and non-radiative carrier recombination. The observed average fluorescence lifetime (⟨*τ*⟩), usually in nanoseconds, depends on radiative (*k*_*R*_) and nonradiative (*k*_*NR*_) relaxation rates as ⟨*τ*⟩ = (*k*_*R*_ + *k*_*NR*_)^−1^ (73). The excited state decay process is also highly sensitive to the local molecular environment due primarily to influences on the available non-radiative carrier relaxation pathways, and thus the non-radiative decay rate, *k*_*NR*_. In principle, lifetime-based FRET can be calculated without any correction factors using equation 17 (9):

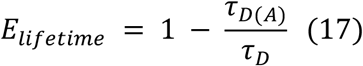

where *τ*_*D*(*A*)_ is the fluorescence lifetime of the donor in the presence of the acceptor, and *τ*_*D*_ is the fluorescence lifetime of the donor in the absence of the acceptor. The fluorescence lifetimes for each benchmark DNA sample were obtained simultaneously with the previous FCS measurements and fit to an exponential function to extract the fluorescence lifetimes, *τ* . Several trends are apparent in the data as shown in Supplementary Figure S1 and summarized in Supplementary Table S4. For the Atto647N-NHS ester dye, *τ*_*Atto*647*N*_ = 3.53 ± 0.00 *ns*, consistent with the value from the manufacturer. Upon attachment to the acceptor DNA strand, *τ*_*Atto*647*N*_ increases to 4.35 ± 0.06 ns, indicating a change in the local environment of the fluorophore likely due to the presence of the DNA nucleotides (73). The finding is similar for the three donor strands. The annealed lo, mid, and hi FRET sample excited state decays required a bi-exponential function to fit the donor data with a 1 – 2 ns component and a 4 ns component. We can attribute the 4 ns component to the decay of the donor molecule in the absence of the acceptor molecule, while the 1 – 2 ns component represents the donor fluorescence influenced by non-radiative energy transfer to the nearby acceptor molecule from FRET. For the faster component, the lo, mid, and hi FRET donor fluorescence lifetimes are 2.13 ± 0.08, 1.37 ± 0.05, 1.19 ± 0.16 ns, respectively. The decreasing lifetime of that relaxation process is consistent with an increasing probability of non-radiative FRET since *τ* = (*k*_*R*_ + *k*_*NR*_)^−1^. FRET efficiency can be calculated from the diffusing lifetime data, but the aggregate results from a sub-ensemble measurement probing thousands of molecules may obscure the calculation, especially if sub-populations are present. If the bursts from individual diffusion events are analyzed, an accurate lifetime-based FRET efficiency may be extracted (24). Using the single-molecule fluorescence intensity traces collected on individual surface-attached DNA duplexes offers a more direct method to calculate lifetime-based FRET efficiency.

To calculate the FRET efficiency for the individual benchmark DNA duplexes depicted in Figure 5, we fit the decays obtained from binning the nanotimes of photons for each signal (*D*_*ex*_*D*_*em*_,*D*_*ex*_*A*_*em*_ (*FRET*) and *A*_*ex*_ *A*_*em*_), and tail-fit the decay to a single exponential function. The excited state decay of the donor signal (*D*_*ex*_ *D*_*em*_) was generated and fit before and after the bleach step to obtain *τ*_*D*(*A*)_and *τ*_*D*_, respectively. The lifetime-based FRET efficiency calculated with equation 17 for the single DNA duplex traces in Figure 5B are – lo FRET: 0.15, mid FRET: 0.52, and hi FRET: 0.66, consistent with the intensity-based FRET efficiency calculation on the same traces, and the values we reported for the diffusing measurements. The values obtained for the three traces in Figure 5B are compared to the diffusing measurements in Table 1. Interestingly, a trend is observed in the *D*_*ex*_ *A*_*em*_ (*FRET*) fluorescence lifetime. This signal represents the fluorescence from the *acceptor* molecule due to energy transfer. With the donor and acceptor farthest apart in the lo sample, the lifetime of the acceptor excited state created from FRET is 5.2 ns, and it decreases to 4.65 and 3.92 for the mid and hi samples, respectively. The same trend in fluorescence lifetimes is not observed for the identical acceptor molecule under direct excitation (*A*_*ex*_ *A*_*em*_), suggesting that the quantum yield of the acceptor is influenced by the distance from the donor molecule in FRET (70). When biomolecules are attached to a surface, a potential concern is introducing artifacts due to surface interactions. We have shown that with careful calibration and determination of correction factors, consistent lifetime and intensity-based FRET efficiency calculations can be obtained from diffusing and surface-attached molecules.

**Table 1:** Fluorescence lifetimes for single-molecule fluorescence intensity traces in Figure 5B. The nanotimes for the *D*_*ex*_*D*_*em*_, *D*_*ex*_*A*_*em*_ (FRET), and *A*_*ex*_*A*_*em*_ signals were separated and tail-fit to a single exponential function. The donor fluorescence emission *D*_*ex*_*D*_*em*_ was further separated into pre- and post-bleach to obtain the fluorescence lifetime of the donor in the presence (*τ*_*D*(*A*)_) and absence (*τ*_*D*_) of the acceptor, respectively. Equation 17 was used to calculate the lifetime-based FRET efficiency. We also calculated the intensity-based FRET efficiency for each single-molecule trajectory, first calculating the gamma (*γ*) factor with equation 16. The average gamma factor determined with the diffusing PIE-FRET measurement was 0.88. The intensity-based FRET efficiency was calculated with equation 14 and the *α* and *δ* correction factors from Supplementary Table S5. (surf. = surface-attached)

| Figure 5 Trace | Direct Acceptor ($A_{ex}A_{em}$) $\tau$ (ns) | FRET ($D_{ex}A_{em}$) $\tau$ (ns) | Donor ($D_{ex}D_{em}$) $\tau$ pre-bleach (ns) | Donor ($D_{ex}D_{em}$) $\tau$ post-bleach (ns) | gamma (surf.) | Lifetime-FRET (surf.) | Intensity FRET (surf.) | Diffusing PIE-FRET |
| --- | --- | --- | --- | --- | --- | --- | --- | --- |
| Lo FRET Trace | 3.63 | 5.2 | 2.81 | 3.3 | 0.74 | 0.15 | 0.11 | 0.15 |
| Mid FRET Trace | 3.77 | 4.65 | 1.74 | 3.66 | 0.59 | 0.52 | 0.54 | 0.49 |
| Hi FRET Trace | 3.7 | 3.92 | 1.08 | 3.21 | 0.64 | 0.66 | 0.68 | 0.76 |

## CONCLUSIONS

Time-resolved fluorescence measurements add several dimensions to optical imaging and spectroscopy for investigation of biomolecules. The ability to measure fluorescence lifetimes, FCS, and PIE-FRET simultaneously on diffusing species leads to a range of possibilities for monitoring transient interactions and conformational dynamics. With the high-resolution photon timing afforded by TCSPC, we can capture dynamics from nanoseconds to seconds, but extracting reliable sample information requires careful calibration. We have presented practical steps to validate experimental FCS and PIE-FRET measurements and applied our calibration to biological systems of interest including liposomes, streptavidin-coated quantum dots, proteins, and nucleic acids. The measured values and trends in the data are consistent with the expected photophysics and relative sizes of the molecules. Experiments conducted on more complex systems may report protein binding, cleavage of nucleic acids by a nuclease, or dynamic protein conformational changes, for example.

Throughout the article, we have compared several methods side-by-side, and each approach has associated limitations and advantages/disadvantages. FCS is a powerful technique, but it is limited to diffusing species. The measurement requires a low sample concentration (usually tens of nanomolar) to ensure only a few molecules are present in the confocal volume at any instant in time, and analysis is hampered if large aggregates are present in solution. The extracted diffusion time, *τ*_*D*_, from a model fit can be converted to a diffusion coefficient, *D*, which requires knowledge of the confocal volume aspect ratio, *κ*. We discussed two methods to obtain *κ*, and the measurement using fluorescent dyes with known diffusion coefficients ultimately achieved more repeatable calculations of *D*.

PIE-FRET isolates diffusing donor-acceptor pairs, and the collected signal results from thousands of molecules. The intensity-based burstwise analysis requires applying correction factors for donor leakage (*α*) and acceptor direct excitation (*δ*), which is necessary for any intensity-based smFRET analysis. The PIE capability offers two approaches to calculate the correction factors – photon counting and Gaussian fitting, which were in very good agreement (Supplementary Table S5). The photon counting method, however, is much more straightforward, requiring only a single analysis step. We achieved excellent agreement with the benchmark study for our calculated FRET efficiency and distance values. Our benchmark samples were a single species that was not dynamic. PIE-FRET analysis becomes challenging and has limitations when multiple subpopulations and dynamic species are in solution. In that case, burstwise fluorescence lifetime-based analysis may be helpful.

Confocal microscopy is also capable of imaging surface-attached single molecules for smFRET studies that are similar to total internal reflection fluorescence (TIRF) microscopy-based smFRET. Importantly, FLIM-FRET provides a direct way to infer energy transfer from an image, thus can easily be extended to live cell experiments. Although diffusing PIE-FRET is relatively high throughput, the residence time of single molecules in the confocal volume is short (∼ 1 ms), and we cannot observe individual complexes for long periods of time. By contrast, monitoring surface-attached single molecules can reveal fast dynamic conformational changes. The speed of acquisition is limited by the laser scan speed (confocal) or camera frame rate (TIRF). We directly compared the FRET efficiency calculated from the diffusing sub-ensemble (PIE-FRET) and single-molecule FLIM-FRET experiments (Table 1). We collected fluorescence intensity traces on single DNA duplexes bound to the coverslip and calculated the FRET efficiency using intensity and lifetime data from the same trace. We applied the *γ* correction factor for the intensity-based calculation, which is more straightforward to determine with surface-attached molecules. The results agree well, although we highlight that the lifetime-based calculation does not require any correction factors. The trend in fluorescence lifetimes also revealed an interesting photophysical effect on the acceptor fluorescence lifetime due to emission from FRET.

We have demonstrated the versatility of TCSPC applied to several biological systems. Using calibrated FCS and PIE-FRET in tandem can reveal conformational and oligomeric states of proteins, nucleic acids, or their complexes for detailed mechanistic studies. Furthermore, studying fast dynamics of surface-attached individual complexes and in live cells with FLIM-FRET is an exciting direction.

## ACRONYMS

A: acceptor
ALEX: alternating laser excitation
APD: avalanche photodiode detector
D: donor
FCS: fluorescence correlation spectroscopy
FCCS: fluorescence cross correlation spectroscopy
FLCS: fluorescence lifetime correlation spectroscopy
FLIM: fluorescence lifetime imaging microscopy
FRET: Förster/fluorescence resonance energy transfer
IPTG: isopropyl β-D-1-thiogalactopyranoside
NA: numerical aperture
PIE: pulsed interleaved excitation
PMT: photomultiplier tube (detector)
smFRET: single-molecule Förster/fluorescence resonance energy transfer
TCSPC: time-correlated single photon counting
TIRF: total internal reflection fluorescence (microscopy)
TTTR: time-tagged time-resolved

## ACKNOWLEDGEMENTS

We thank PicoQuant for remote installation, training, and support. We especially thank Dr. Olaf Schulz, Dr. Steffen Ruettinger, Dr. Evangelos Sisamakis, and Dr. Samaneh Rezvani. We thank Dr. Robin Stanley and lab at the National Institute of Environment Health Sciences (NIEHS) - Dr. Meredith Frazer, Isha Wilson, and Dr. Zoe Wright – for Nsp15 expression plasmids and purification support. We thank Dr. Keith Weninger for critical reading of the manuscript and helpful discussions. We thank Dr. Dorothy Erie and Dr. Manju Hingorani for MutL expression plasmids.

## AUTHOR CONTRIBUTIONS

S.J.L. designed the experiments, analyzed and interpreted data, and prepared the manuscript. B.S.C. conducted FCS calibration experiments and data analysis. I.S. conducted PIE-FRET calibration experiments with benchmark DNA substrates and data analysis. K.G. conducted PIE-FRET experiments and data analysis for the RNA-DNA hybrid substrates and prepared and labeled the Nsp15 protein. J.F.C. prepared and labeled the MutL protein, conducted diffusion experiments and performed PAM analysis. L.A.R. conducted the single molecule FLIM-FRET experiments with the benchmark DNA substrates. A.N.M. assisted with the technical aspects of all experiments.

## DATA AVAILABILITY

SymPhoTime files (.ptu) are available upon request. Open-source PIE analysis with MATLAB (PAM) software (https://gitlab.com/PAM-PIE) can open and process .ptu files.

