## Supplementary Information for "A practical guide to time-resolved fluorescence microscopy and spectroscopy"

#### I. Sequences for benchmark DNA substrates

**Supplementary Table S1:** Sequences for the acceptor (A) and donor (D) strands that were purchased with a 6-carbon linker-$NH_2$-deoxythymidine (internal amino modifier C6 dT) modification for fluorescent labeling via N-hydroxysuccinimide ester (NHS-ester) attachment chemistry. The double underline indicates the amino-dT modification on the acceptor and donor strands. The acceptor strand also contains a 5'-biotin modification for surface attachment. The strands are separately labeled with an acceptor dye (Atto647N-NHS ester) or donor dye (Atto550-NHS ester). The same labeled A strand is used for all constructs. Annealing of the A strand with one of the D strands results in duplex DNA with the indicated base pair (bp) separation for low (lo), middle (mid), or high (hi) FRET signal.

| Oligo | Amino-dT Position (Linker) | Sequence (5' – 3') | Annealed D-A Separation |
|---|---|---|---|
| Acceptor strand | T31(C6) | 5' - biotin - CCA GAC AAA CAC TCA AAC AAA CTC GAC ACT TTC AGC TC - 3' | n/a |
| Donor strand (Lo FRET) | T31(C6) | 5' - GAG CTG AAA GTG TCG AGT TTG TTT GAG TGT TTG TCT GG - 3' | 23 bp |
| Donor strand (Mid FRET) | T23(C6) | 5' - GAG CTG AAA GTG TCG AGT TTG TTT GAG TGT TTG TCT GG - 3' | 15 bp |
| Donor strand (Hi FRET) | T19(C6) | 5' - GAG CTG AAA GTG TCG AGT TTG TTT GAG TGT TTG TCT GG - 3' | 11 bp |

#### II. Calculation of labeling efficiency for benchmark DNA oligomers

We fluorescently labeled the benchmark DNA oligomers listed in Supplementary Table S1 with either a donor or acceptor dye as described in the main text (Materials and Methods). To determine the labeling efficiency, we measured the absorbance (*A, Abs*) on a UV/Vis spectrophotometer (Agilent Cary 3500). We first measured the absorbance spectrum of the free dyes in water at a micromolar concentration. The absorbance of the dyes at 260 nm and the peak absorbance of the dye listed in the table below were used as correction factors.

| Sample | Abs. at 260 nm | Abs. at 554 nm | Abs. at 646 nm |
|---|---|---|---|
| Atto550 NHS-Ester | 0.0651 | 0.2807 | n/a |
| Atto647N NHS-Ester | 0.0276 | n/a | 0.71 |

The correction factors are calculated using this ratio:

$$CF = \frac{A_{dye}(260\,nm)}{A_{dye}(\lambda_{dye})}$$

, where $A_{dye}(260\,nm)$ is the absorbance of the dye at the wavelength at 260 nm, and $A_{dye}(\lambda_{dye})$ is the absorbance of the dye (either 550 nm or 647 nm). The correction factors (*CF*) we calculated for the two dyes are 0.232 for 550 nm and 0.039 for 647 nm. We measured the absorbance spectrum of each fluorescently labeled oligo sample and corrected the oligo absorbance at 260 nm using the following equation:

$$A_{oligo} = A_{oligo}(260\,nm) - \left(A_{oligo}(550\,nm) \cdot CF_{550\,nm}\right) - \left(A_{oligo}(647\,nm) \cdot CF_{647}\right)$$

The dye and oligo concentrations were calculated using the Beer-Lambert law with a calculated value for the dye and oligo extinction coefficients ($\varepsilon$) in $\frac{L}{mol \cdot cm}$ (values obtained from the manufacturers).

$$Concentration_{dye} = \frac{A_{dye}(\lambda_{dye})}{\varepsilon_{dye}}$$

$$Concentration_{oligo} = \frac{A_{oligo}}{\varepsilon_{oligo}}$$

The oligo labeling efficiency (*E*) was calculated using the following ratio:

$$E = \frac{Concentration_{dye}}{Concentration_{oligo}}$$

Supplementary Table S2 summarizes the absorbance values, calculated concentrations, and labeling efficiencies for each fluorescently labeled DNA strand. The values of $\varepsilon$ were reported in the manufacturer specification sheet (Integrated DNA Technologies), and are listed below:

Acceptor: $\varepsilon$ = 364,500 L/mol-cm
Lo donor: $\varepsilon$ = 366,300 L/mol-cm
Mid donor: $\varepsilon$ = 366,300 L/mol-cm
Hi donor: $\varepsilon$ = 366,300 L/mol-cm

**Supplementary Table S2:** UV/Vis absorbance measurements and oligo labeling efficiencies. A percentage above 100 may indicate the presence of small amount free dye.

| | Absorbance at Wavelength (nm) | | | | | | | | |
|---|---|---|---|---|---|---|---|---|---|
| **Sample (dilution ratio)** | **260** | **550** | **647** | **Adjusted Abs. at 260 nm** | **Atto550 Conc. (M)** | **Atto647N Conc. (M)** | **Oligo Conc. (M)** | **Atto550 Labeling Efficiency** | **Atto647N Labeling Efficiency** |
| Acceptor (1:50) | 0.70 | n/a | 0.34 | 0.69 | n/a | 2.24E-06 | 1.89E-06 | 0.00% | 118.30% |
| Lo Donor (1:100) | 0.14 | 0.05 | n/a | 0.13 | 3.88E-07 | n/a | 3.49E-07 | 111.14% | n/a |
| Mid Donor (1:100) | 0.17 | 0.061 | n/a | 0.16 | 5.07E-07 | n/a | 4.33E-07 | 116.95% | n/a |
| Hi Donor (1:100) | 0.18 | 0.032 | n/a | 0.17 | 2.70E-07 | n/a | 4.72E-07 | 57.24% | n/a |
| Annealed Lo (1:100) | 0.012 | 0.002 | 0.0024 | 0.012 | 1.58E-08 | 1.60E-08 | 3.17E-08 | 49.90% | 50.42% |
| Annealed Mid (1:50) | 0.032 | 0.009 | 0.006 | 0.03 | 7.50E-08 | 4.13E-08 | 8.03E-08 | 93.39% | 51.47% |
| Annealed Hi (1:100) | 0.014 | 0.0011 | 0.005 | 0.014 | 9.17E-09 | 3.07E-08 | 3.83E-08 | 23.92% | 80.04% |

**Supplementary Table S3:** Summary of the confocal volume parameters determined from imaging 100 nm Tetraspek® beads on a glass coverslip (top) and from fitting FCS curves of organic dyes with known diffusion coefficients. Diffusion coefficients for the dyes were obtained from published values (1). The reported errors for the bead imaging values are calculated from the covariance of the 2D elliptical Gaussian fits. The reported error for each curve fitting parameter is a combination of statistical and fit covariance error, $\sigma_{total} = \sqrt{\sigma_{covar}^2 + \sigma_{stats}^2}$.

| **Laser wavelength (nanometers, nm)** | Effective volume, $V_{eff}$ **(femtoliters, fL)** | **Aspect ratio,** $\kappa$ | **Lateral radius,** $\omega_0$ **(nanometers, nm)** |
|---|---|---|---|
| *Bead imaging* | | | |
| 485 | 0.823 ± 0.005 | 3.343 ± 0.007 | 353 ± 0 |
| 531 | 0.981 ± 0.006 | 3.395 ± 0.007 | 373 ± 0 |
| 636 | 1.619 ± 0.010 | 3.037 ± 0.006 | 457 ± 0 |
| *Curve fitting FCS measurements of organic dyes* | | | |
| 485 (Fluorescein, Rhodamine 110, Rhodamine 123) | 1.35 ± 0.08 | 8.3 ± 0.7 | |
| 531 (Rhodamine B, Rhodamine 6G) | 1.54 ± 0.10 | 7.1 ± 0.8 | |
| 636 (Atto655 maleimide) | 1.75 ± 0.06 | 3.8 ± 0.5 | |

#### III. Fluorescence and diffusion characteristics of free dyes and labeled benchmark DNA

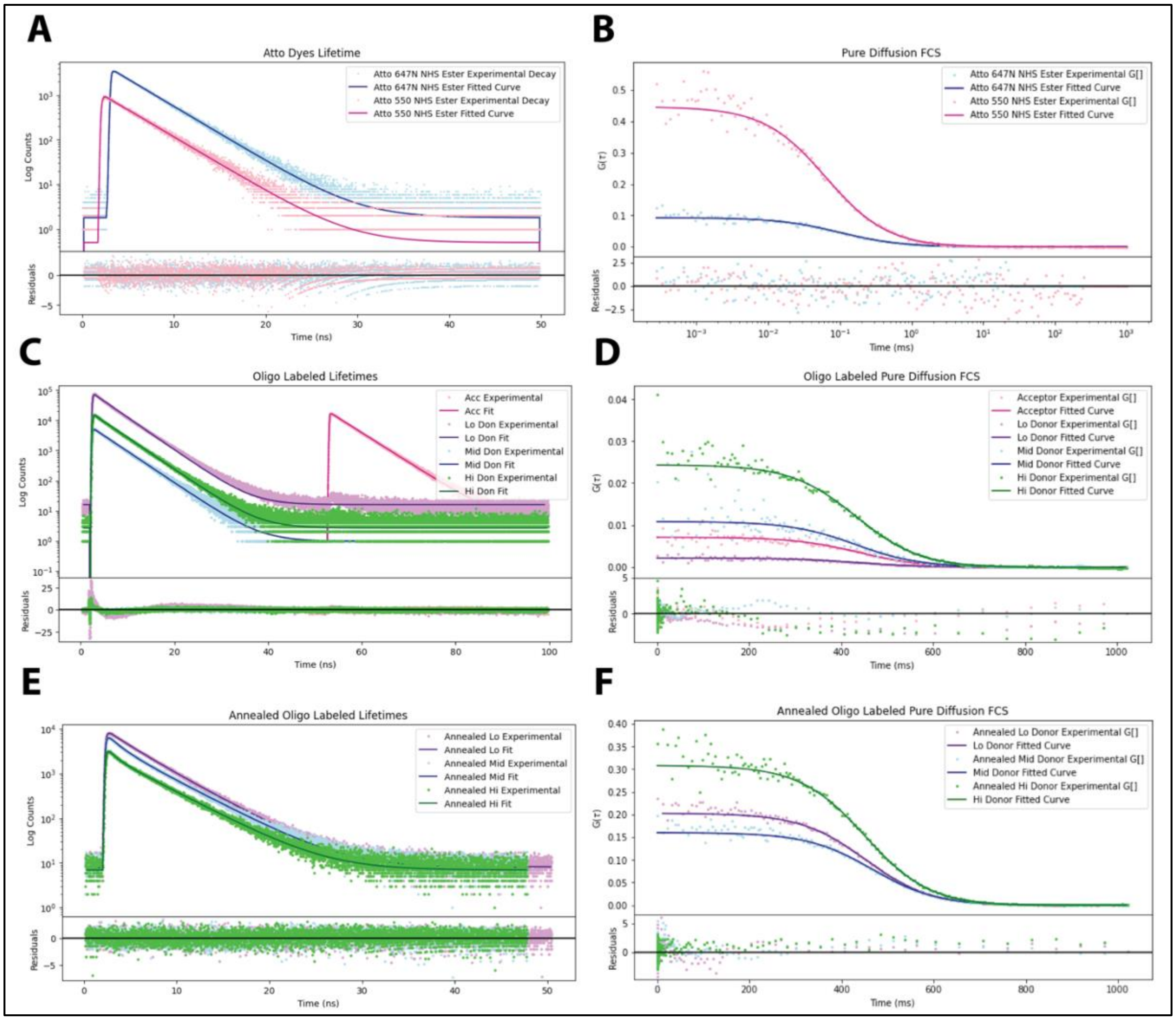


**Supplementary Figure S1:** Representative fluorescence lifetimes and diffusion coefficients for free Atto 550 and Atto 647N NHS ester dyes, labeled single donor (D) and acceptor (A) benchmark DNA strands (Table S1), and annealed D-A substrates (all in PBS). Experimental excited state decays (dots) were fit to a single or double exponential function by iterative reconvolution with an instrument response function in the SymPhoTime software (black line) and the fluorescence lifetimes ($\tau$) were extracted (**A, C, E**). An autocorrelation was performed on the fluorescence intensity time trace and the autocorrelation decays (dots) were fit to a pure diffusion model (black lines, see main text equation 2) to extract diffusion coefficients **(B, D, F)**. **A)** The representative fluorescence lifetimes of free dyes were $\tau$ = 3.60 ns ($\chi^2 = 1.05$) and $\tau$ = 3.53 ns ($\chi^2 = 1.04$) for Atto550 and Atto647N, respectively. **B)** The representative diffusion coefficients, $D$, from the fits were 352 $\frac{\mu m^2}{s}$ ($\chi^2$ = 1.48) and 323 $\frac{\mu m^2}{s}$ ($\chi^2 = 1.36$) for Atto 550 and Atto 647N, respectively. **C)** The fluorescence lifetimes for the single donor and acceptor strands are 4.43 ns ($\chi^2 = 1.62$), 4.05 ns ($\chi^2 = 11.86$), 4.19 ns ($\chi^2 = 1.87$), and 4.14 ns ($\chi^2 = 2.62$), for acceptor, lo donor, mid donor, and hi donor strands, respectively. **D)** The diffusion coefficients, $D$, for the single DNA strands were 121 $\frac{\mu m^2}{s}$ ($\chi^2$ = 0.83), 125 $\frac{\mu m^2}{s}$ ($\chi^2$ = 1.14), 132 $\frac{\mu m^2}{s}$ ($\chi^2$ = 0.75), and 130 $\frac{\mu m^2}{s}$ ($\chi^2$ = 1.82) for acceptor, lo donor, mid donor, and hi donor strands, respectively. **E)** The fluorescence decays for labeled annealed lo FRET, mid FRET, and hi FRET substrates were fit to a sum of two exponentials and the values were $\tau_1$= 2.33 ns and $\tau_2$= 4 ns ($\chi^2 = 1.0$) ; $\tau_1$ =1.32 ns and $\tau_2$ = 4.04 ns ($\chi^2 = 1.01$) ; $\tau_1$= 0.94 ns and $\tau_2$ = 4.05 ns ($\chi^2 = 1.03$) for lo FRET, mid FRET, and hi FRET substrates, respectively. **F)** The diffusion coefficients, $D$, for the annealed substrates were 99 $\frac{\mu m^2}{s}$ ($\chi^2 =$ 2.15), 80 $\frac{\mu m^2}{s}$ ($\chi^2 = 2.17$), and 100 $\frac{\mu m^2}{s}$ ($\chi^2 = 1.68$) for lo FRET, mid FRET, and hi FRET substrates, respectively. The average values for each sample are summarized in Supplementary Table S4 below.

**Supplementary Table S4:** Summary of fluorescence lifetime and diffusion coefficients for benchmark DNA substrates (see Table S1 for sequences). Supplementary Figure S1 shows representative decays for each free dye, fluorescently labeled single strand, and annealed sample. The average values from at least three 3-minute measurements are reported in the table below, representing thousands of molecules. For the annealed samples, only the donor lifetime (donor excitation, Ch1 + Ch2 signal - $D_{ex}D_{em} + D_{ex}A_{em}$) is reported, and the relative contribution calculated from the amplitudes of bi-exponential fits are indicated in parentheses. The reported diffusion coefficients are obtained from correlating and fitting the FRET signal only (donor excitation, Ch1 signal – $D_{ex}A_{em}$) to a conformational model (equation 3 in the main text).

| **Sample** | **Lifetime, $\tau$ (ns) (Atto550 donor)** | **Diffusion coefficient, $D$ ($\frac{\mu m^2}{s}$)** |
|---|---|---|
| Atto550 NHS ester | 3.58 ± 0.05 | 342 ± 7 |
| Atto647N NHS ester | 3.53 ± 0.00 | 328 ± 13 |
| Acceptor strand | 4.35 ± 0.06 | 97 ± 19 |
| Donor strand (Lo FRET) | 4.05 ± 0.00 | 150 ± 32 |
| Donor strand (Mid FRET) | 4.12 ± 0.07 | 130 ± 40 |
| Donor strand (Hi FRET) | 4.14 ± 0.01 | 170 ± 26 |
| Annealed Lo FRET | $\tau_1$= 2.25 ± 0.21 (33 %)<br>$\tau_2$= 4.06 ± 0.07 (67 %) | 91 ± 22 |
| Annealed Mid FRET | $\tau_1$= 1.46 ± 0.07 (31 %)<br>$\tau_2$ = 4.24 ± 0.01 (69 %) | 85 ± 5 |
| Annealed Hi FRET | $\tau_1$= 1.04 ± 0.05 (31 %)<br>$\tau_2$ = 4.02 ± 0.03 (69 %) | 108 ± 4 |

### IV. PIE-FRET calibration

**Supplementary Table S5:** PIE-FRET $\alpha$ and $\delta$ calibration values determined by two methods. For the photon count method with fluorescence lifetime data, the reported average and standard deviation is from at least three 3-minute measurements on the same sample. For the Gaussian fitting method with intensity data, the reported error is the average of the Gaussian peak value, $b_1$, from at least two 2-minute measurements representing thousands of molecules.

| **Correction Factors** | **Calculated using photon counts** | **Calculated using fitted Gaussian curve (peak)** |
|---|---|---|
| **Spectral crosstalk** | α | α |
| Atto550 free dye | 0.069 ± 0.001 | 0.069 |
| Donor Lo – Atto550 | 0.061 ± 0.000 | 0.053 |
| Donor Mid – Atto550 | 0.065 ± 0.002 | 0.060 |
| Donor Hi – Atto550 | 0.061 ± 0.000 | 0.054 |
| AF546 maleimide | 0.059 ± 0.001 | 0.058 |
| AF546-DNA complement to RNA/DNA hybrid | 0.058 ± 0.002 | 0.044 ± 0.003 |
| **Direct excitation** | δ | δ |
| Atto647N free dye | 0.029 ± 0.002 | 0.03 |
| Acceptor-Atto647N | 0.021 ± 0.000 | 0.019 |
| Atto647N-RNA/DNA hybrid | 0.030 ± 0.003 | 0.033 ± 0.003 |

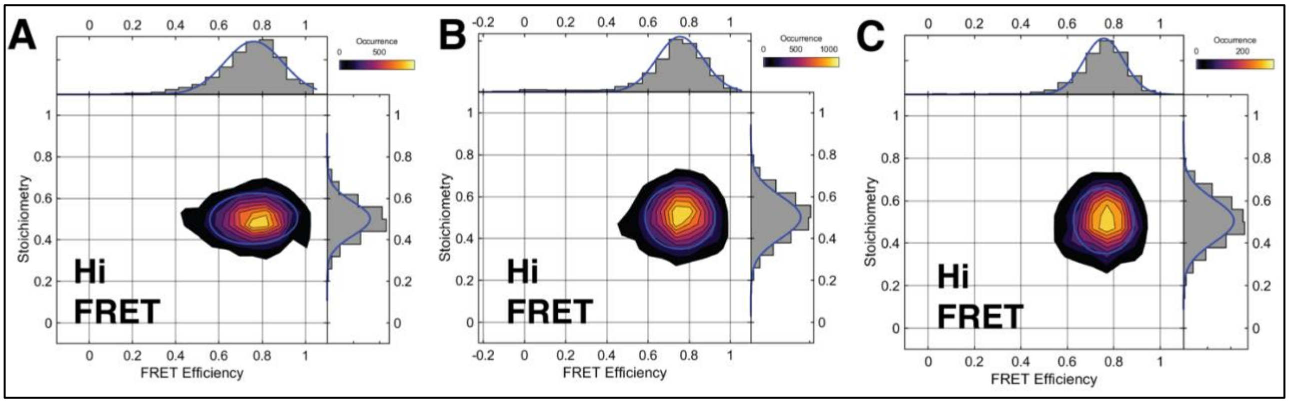


**Supplementary Figure S2:** Comparison of different concentration and acquisition times for the Hi FRET sample. The parameters for each measurement extracted from analysis in the PAM software are summarized below. A longer acquisition time at low DNA concentration did result in a narrower distribution, but the same mean FRET efficiency, $\langle E \rangle$, was determined for each measurement. The Hi FRET A plot is the result of aggregated data from three 3-minute measurements. *DCBS is dual channel burst search, and APBS is all-photon burst search that was used in conjunction with an ALEX-2CDE filter, removing donor only, acceptor only, and mixture subpopulations (2). All data were corrected for $\alpha, \delta, \gamma$ and $\beta$ as described in the main text.

| Sample | Parameters | | | | | | | |
|---|---|---|---|---|---|---|---|---|
| | **Conc. (nM)** | **Laser power (μW)** | **Acq. Time** | **Burst algorithm** | **Mean FRET** | **$\sigma$** FRET | **$\gamma$** | **$\beta$** |
| Hi FRET A | 20 | 2.3 | 3 min. (x3) | DCBS | 0.76 | 0.13 | 0.88 | 1.05 |
| Hi FRET B | 2 | 9.1 | 3 hours | DCBS | 0.76 | 0.11 | 0.88 | 1.2 |
| Hi FRET C | 2 | 9.1 | 3 hours | APBS | 0.76 | 0.09 | 0.88 | 1.4 |

**Supplementary Table S6:** Gaussian fit parameters for PIE-FRET histograms in Figure 4 in the main text. We fit the histograms to a Gaussian function using the PAM software.

| Sample | Fitting parameters | | | | | |
|---|---|---|---|---|---|---|
| | **Mean FRET, $\langle E \rangle$** | **$\sigma$ FRET** | **Mean Stoich.** | **$\sigma$ Stoich.** | **$\gamma$** | **$\beta$** |
| Lo FRET post correction | 0.15 | 0.11 | 0.48 | 0.08 | 0.88 | 0.52 |
| Mid FRET post correction | 0.49 | 0.16 | 0.49 | 0.09 | 0.88 | 0.52 |
| Hi FRET post correction | 0.76 | 0.13 | 0.50 | 0.08 | 0.88 | 1.05 |

The calculated FRET efficiency can be converted to a physical distance, $R$, by rearranging the theoretical FRET efficiency expression ($E_{FRET} = [1 + (R/R_0)^6]^{-1}$) to obtain the following equation:

$$R = R_0 \left[ \frac{1}{E} - 1 \right]^{1/6}$$

, where $R$ is the donor-acceptor separation, $R_0$ is the Förster radius, and $E$ is the FRET efficiency calculated with equation 14 in the main text. The distance histograms were calculated on a burst wise intensity basis for each lo, mid, and hi FRET sample pre- and post-correction, and the results are presented in Supplementary Figure S3. The x-axis for the distance histograms is in units of $R_0$, which is the distance at which FRET efficiency is 50% for any given FRET pair.

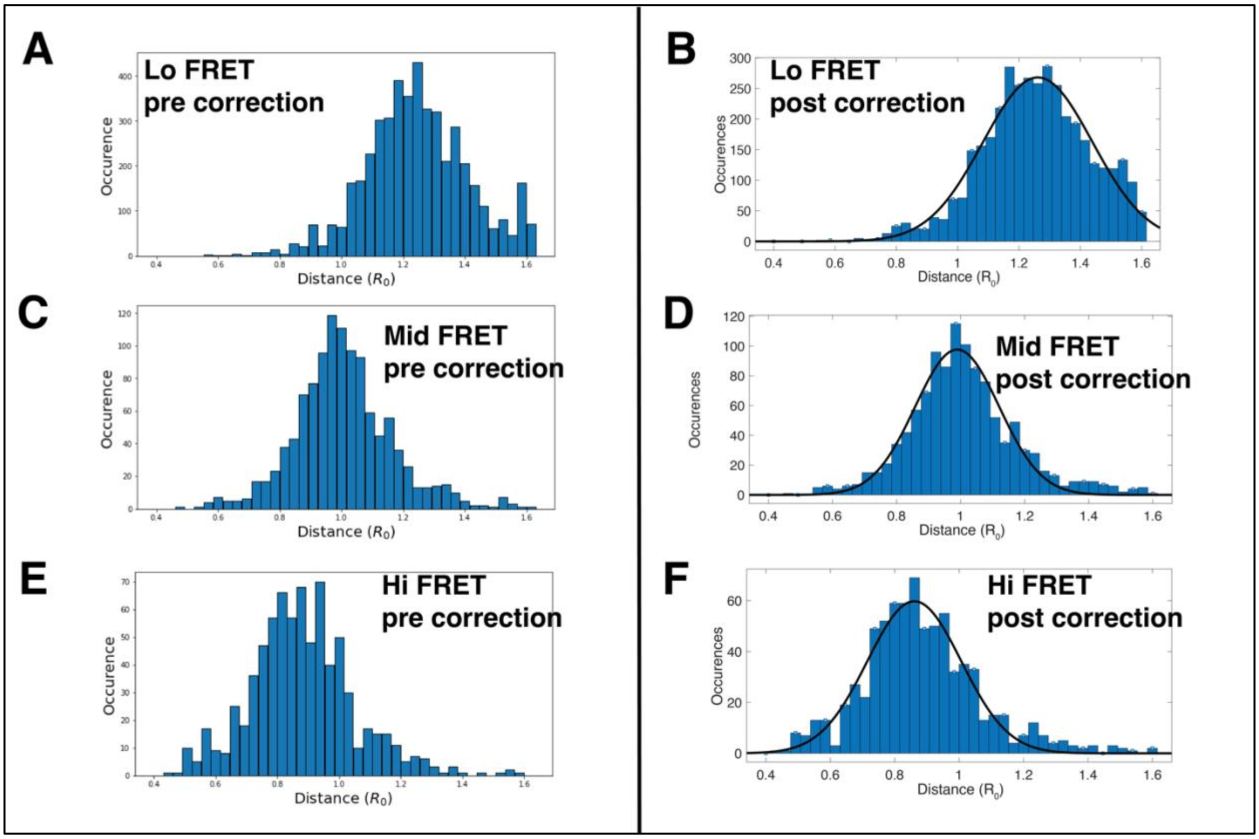


**Supplementary Figure S3:** Distance histograms for lo, mid, and hi FRET substrates presented in Figure 4 of the main text. The post-correction histograms (B, D, F) were fit to a Gaussian function of the form $f(x) = a_1 * e^{(-((x-b_1)/c_1)^2)}$, and the peak position (b1) was used as the average distance between Atto dyes attached to the benchmark DNA. From the previous benchmark study(3), $R_0$ for the Atto550 – Atto647N dye pair was determined to be 6.26 nm, thus the donor-acceptor distance can be obtained from multiplying the x-axis values by 6.26 nm. The physical distance between label sites was calculated by assuming 0.34 nm per base pair. The table below summarizes the calculated physical distance between label sites and the average dye separation in each annealed DNA samples determined from fitting the distance histograms above.

| **Sample** | **Base pair separation** | **Physical distance (nm)** | **Distance from Gaussian fit ($R_0$ units)** | **Average distance between dyes (nm)** |
|---|---|---|---|---|
| Lo FRET, post correction | 23 | 7.82 | 1.26 | 7.89 |
| Mid FRET, post correction | 15 | 5.1 | 0.99 | 6.2 |
| Hi FRET, post correction | 11 | 3.74 | 0.86 | 5.38 |

#### V. RNA/DNA hybrid substrates

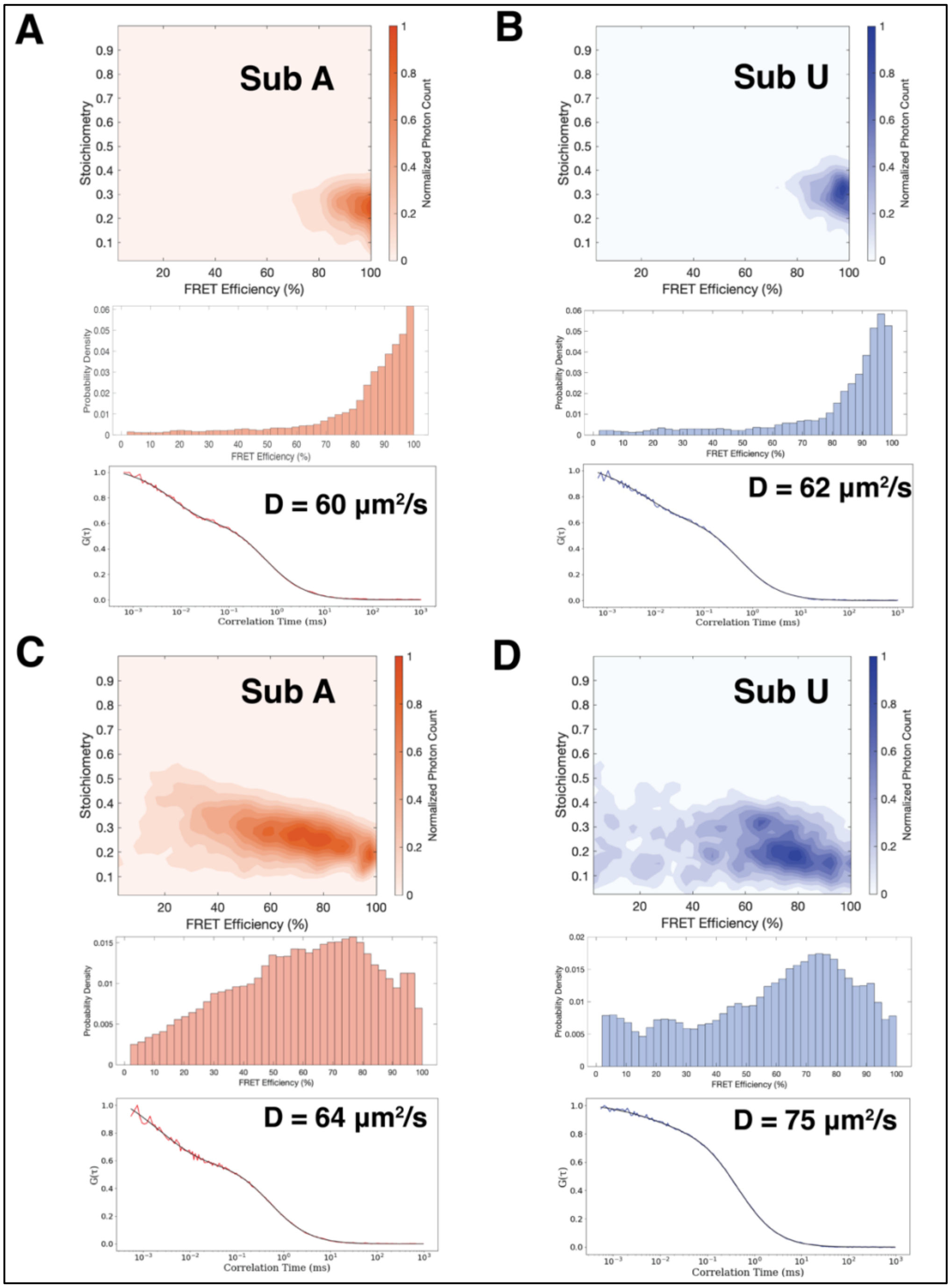


**Supplementary Figure S4:** Two-dimensional *E-S* histograms, 1-D PIE-FRET histograms, and FCS curves with conformational fits of hybrid RNA-DNA substrates (see the Materials and Methods section in the main text for sequences). The PIE-FRET histograms were corrected with the $\alpha$ and $\delta$ values from Supplementary Table S5.

**A)** and **B)** were collected with the 10-base D-A separation construct, while **C)** and **D)** were collected with the 15-base D-A separation construct.
